# The relation between crosstalk and gene regulation form revisited

**DOI:** 10.1101/372672

**Authors:** Rok Grah, Tamar Friedlander

**Affiliations:** Institute of Science and Technology Austria, Am Campus 1, A-3400 Klosterneuburg, Austria; The Robert H. Smith Institute of Plant Sciences and Genetics in Agriculture Faculty of Agriculture, Hebrew University of Jerusalem, P.O. Box 12 Rehovot 7610001, Israel

## Abstract

Genes differ in the frequency at which they are expressed and in the form of regulation used to control their activity. In particular, positive or negative regulation can lead to activation of a gene in response to an external signal. Previous works proposed that the form of regulation of a gene correlates with its frequency of usage: positive regulation when the gene is frequently expressed and negative regulation when infrequently expressed. Such network design means that, in the absence of their regulators, the genes are found in their least required activity state, hence regulatory intervention is often necessary. Due to the multitude of genes and regulators, spurious binding and unbinding events, called “crosstalk”, could occur. To determine how the form of regulation affects the global crosstalk in the network, we used a mathematical model that includes multiple regulators and multiple target genes. We found that crosstalk depends non-monotonically on the availability of regulators. Our analysis showed that excess use of regulation entailed by the formerly suggested network design caused high crosstalk levels in a large part of the parameter space. We therefore considered the opposite ‘idle’ design, where the default unregulated state of genes is their frequently required activity state. We found, that ‘idle’ design minimized the use of regulation and thus minimized crosstalk. In addition, we estimated global crosstalk of *S. cerevisiae* using transcription factors binding data. We demonstrated that even partial network data could suffice to estimate its global crosstalk, suggesting its applicability to additional organisms. We found that *S. cerevisiae* estimated crosstalk is lower than that of a random network, suggesting that natural selection reduces crosstalk. In summary, our study highlights a new type of protein production cost which is typically overlooked: that of regulatory interference caused by the presence of excess regulators in the cell. It demonstrates the importance of whole-network descriptions, which could show effects missed by single-gene models.

**Author Summary:** Genes differ in the frequency at which they are expressed and in the form of regulation used to control their activity. The basic level of regulation is mediated by different types of DNA-binding proteins, where each type regulates particular gene(s). We distinguish between two basic forms of regulation: positive – if a gene is activated by the binding of its regulatory protein, and negative – if it is active unless bound by its regulatory protein. Due to the multitude of genes and regulators, spurious binding and unbinding events, called “crosstalk”, could occur. How does the form of regulation, positive or negative, affect the extent of regulatory crosstalk? To address this question, we used a mathematical model integrating many genes and many regulators. As intuition suggests, we found that in most of the parameter space, crosstalk increased with the availability of regulators. We propose, that crosstalk is usually reduced when networks are designed such that minimal regulation is needed, which we call the ‘idle’ design. In other words: a frequently needed gene will use negative regulation and conversely, a scarcely needed gene will employ positive regulation. In both cases, the requirement for the regulators is minimized. In addition, we demonstrate how crosstalk can be calculated from available datasets and discuss the technical challenges in such calculation, specifically data incompleteness.

## Introduction

Gene regulatory networks can employ different architectures that seemingly realize the same input-output relation. There is a basic dichotomy of gene regulation into positive and negative control. A gene controlled by positive regulation is, by default, not expressed and requires binding of an activator to its operator to induce it. In contrast, a gene controlled by negative regulation, is expressed by default, unless a repressor binds its operator and attenuates its activity. While a gene can be regulated using either mode, researchers have pondered whether additional considerations could favor the choice of one mechanism over the other, or whether this choice is merely a coincidence (“evolutionary accident”). Through-out the years, this question was addressed using different approaches. The seminal work of Michael Savageau [1, 2, 3] proposed the so-called “Savageau demand rule”, namely, that genes encoding frequently needed products (“high-demand”) are often regulated by activators. Conversely, genes whose products are only needed sporadically (“low-demand”), tend to be regulated by repressors. Savageau argued that the intensity of selection depends on the extent to which the regulatory construct is used (later called the “use it or lose it” principle [4]). When infrequently used (as in activator regulating a low-demand or a repressor regulating a high-demand gene), selection to preserve is weak, rendering it unlikely to survive [5]. A later evolutionary analysis mathematically formulated the problem as selection in an alternating environment and found the exact conditions under which the Savageau demand rule is expected to hold [4].

Recently, a comprehensive survey of regulatory topologies in *E. coli* and *B. subtilis*, found agreement between the experimentally observed topologies and their satisfaction of dynamic constraints, as verified in simulations. In light of these exceptions to the Savageau demand rule the authors proposed that evolutionary processes randomly pick a regulatory topology out of the many possible ones meeting the organism physiological constraints [6].

An alternative reasoning for the observed correlation between a gene’s demand and its form of regulation was proposed using a biophysical, rather than evolutionary argument [7, 8]. If a high-demand gene is regulated by an activator and a low-demand gene is regulated by a repressor, their regulatory binding sites are mostly occupied and protected from spurious binding of foreign regulators that could interfere with the gene’s regulatory state. However, if this reasoning applies not just to one gene, but to many of them, it would also entail extravagant use of regulators [6]. This would place heavy demands on protein expression systems, associated with reduced growth rate [9, 10, 11, 12, 13].

While the above-mentioned studies examined the significance of regulatory architectures from different perspectives, they all concentrated on a single gene with a single regulator, regardless of the full regulatory network. It remains unanswered whether the choice of positive or negative regulation for a gene with low- or high-demand could have additional costs for the entire network. Specifically, transcription factors are known to have limited specificity and bind a variety of DNA targets, besides their cognate binding sites [14, 15, 16, 17, 18]. The probability of such binding events naturally depends on their concentrations [19, 20]. Here, we revisit the argument that the Savageau demand rule minimizes transcriptional crosstalk, by accounting for crosstalk of multiple genes simultaneously, rather than the single-gene crosstalk considered earlier.

We use a mathematical global crosstalk model [21], which was built upon the well-established thermodynamic model of gene regulation to calculate transcription factor (TF)-DNA interactions [22, 23, 19, 20, 24, 25, 26]. We have previously shown that while crosstalk affecting a particular gene can be reduced by different means, it always comes at the cost of elevating crosstalk in other genes. In contrast, the *global* crosstalk cannot be reduced below a certain threshold. Here, we analyze global crosstalk levels under different regulatory strategies: either positive or negative regulation. We compare two extreme designs: a ‘busy’ one that implements the Savageau demand rule, in which a high (low)-demand gene is always regulated by an activator (repressor) and an opposite ‘idle’ design, in which a high (low)-demand gene is always regulated by a repressor (activator). We find that the ‘busy’ design maximizes regulator usage, whereas the ‘idle’ one minimizes it. We analyze the dependence of global crosstalk on the abundance of regulatory proteins in the cellular environment and find the exact conditions under which either ‘idle’ or ‘busy’ design minimizes crosstalk. We conclude that under most biologically plausible parameter values, the ‘idle’ design should yield lower *global* transcriptional crosstalk.

This paper begins with the introduction of a general symmetric model for the analysis of transcriptional crosstalk in a many-TFs-many-genes setting, with combination of positive and negative regulation. We show that global crosstalk levels directly depend on the fraction of TFs in use and only indirectly on the choice of activation or repression as the form of regulation. We then analyze TF usage and crosstalk levels of the two extreme designs, i.e., ‘busy’ and ‘idle’ and then construct numerical simulations of a more general asymmetric gene usage model, that are in agreement with the analytical result. Lastly, we discuss the challenges in crosstalk calculation for real gene regulatory networks, in particular, the possible effect of data incompleteness, and show an example using *S. cerevisiae* TF data.

## Results

### A model of gene regulation using a combination of activators and repressors

We consider a cell that has a total of *M* genes, each of which is transcriptionally regulated to be either active or inactive. We assume that each gene is regulated by a single unique TF species - its cognate one. Each gene has a short DNA binding site to which its cognate TF binds. A fraction 0 ≤ *p* ≤ 1 of the genes is regulated by activators and the remaining 1 − *p* fraction of genes is regulated by repressors. When no activator is bound, activator-regulated genes are inactive (or active at a low basal level) and only become active once an activator TF binds their binding site. In contrast, repressor-regulated genes are active, unless a repressor TF binds their binding site and inhibits their activity (Fig 1A). We assume that different environmental conditions require the activity of different subsets of the *M* genes. We assume however that all these subsets include the same proportion of genes 0 ≤ *q* ≤ 1 that is needed to be active. The remaining 1 − *q* proportion should be inactive. These activity states are regulated by the binding and unbinding of the TFs specialized for these genes. We assume that only a subset of TFs necessary to maintain the desired regulatory pattern, are available to bind and regulate these genes. However, TFs often have limited specificity to their DNA targets and can occasionally bind slightly different sequences, albeit with lower probability [27, 16, 17, 28, 29].

**Figure 1:**
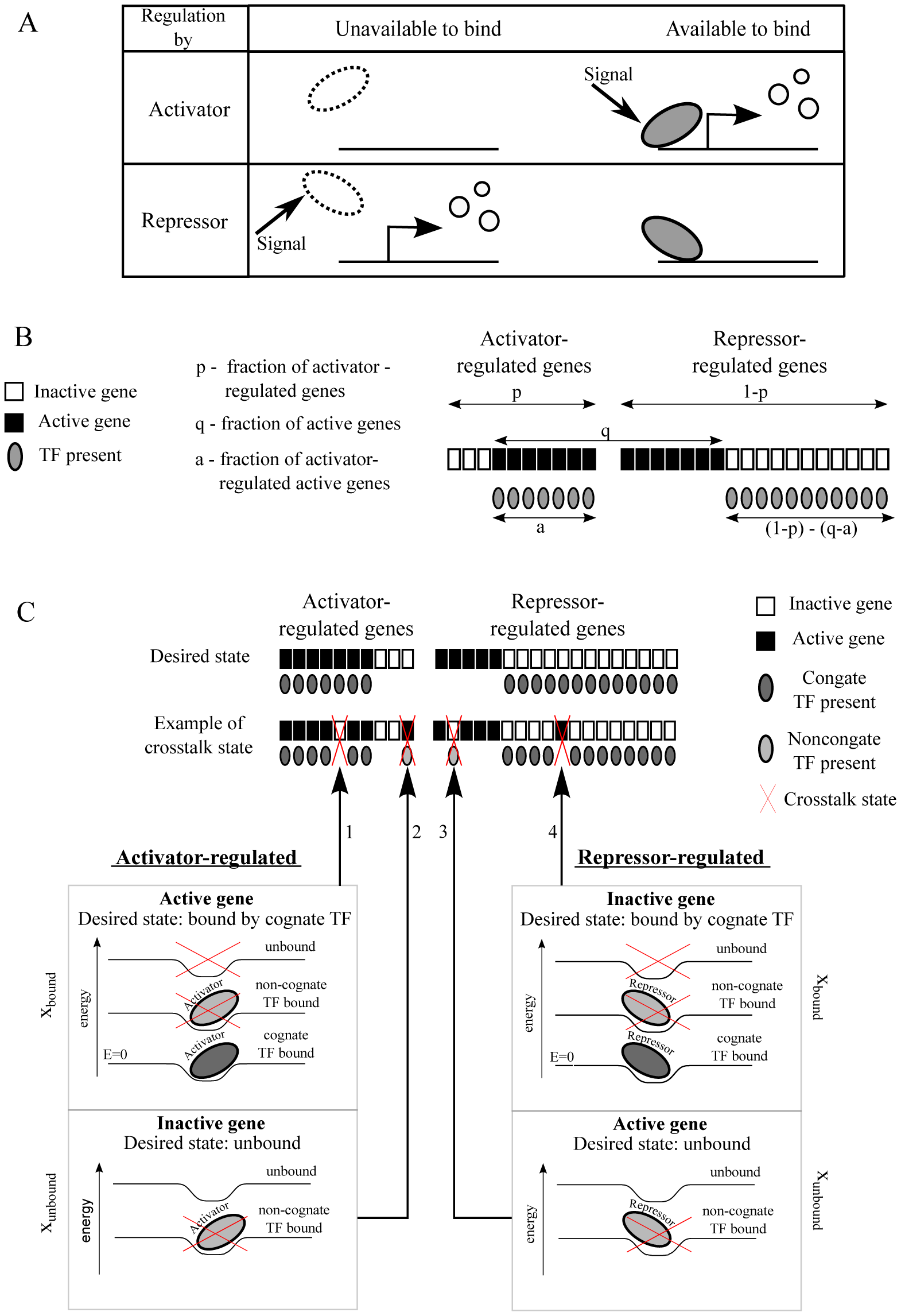
Gene regulation can employ different combinations of activators and repressors to implement the same gene expression pattern. **(A)** A signal can cause gene activation by either positive (first row) or negative (second row) control. **(B)** We consider a total of *M* genes in a cell, of which fraction 0 ≤ *p* ≤ 1 is regulated by activators, and the remaining 1 − *p* is regulated by repressors. Assume that only fraction *q* <1 of these genes should be active under certain conditions (black squares), while the remaining genes should be inactive (white squares). In general, *a* ≤ *q, p* of this *q* proportion is activator-regulated and *q* − *a* is repressor-regulated. Here, we illustrate all four cases of active/inactive genes regulated by activator/repressor and define all the variables. Gray ellipses represent TFs (of either type) required to maintain the regulatory state of the genes. **(C)** Different genes are regulated by different TF species, where TF specificity is determined by short regulatory DNA sequences (binding sites) adjacent to the gene. Each such binding site can be at different levels of energy depending on its TF occupancy. It is in the lowest *E* = 0 (most favorable) level when bound by its cognate TF; it can be in a variety of higher energy levels if a non-cognate TF binds or if the site remains unoccupied (lower panel). The upper panel shows the crosstalk-free ‘desired state’ (first row), where each TF binds its cognate target. Below (second row), four different possibilities in which binding of a TF to non-cognate binding sites or failure to bind lead to crosstalk. An activator-regulated gene should ideally be regulated by its cognate activator (right-inclined ellipse), in order to become active. If this cognate TF fails to bind when the gene should be active (1), or if another TF binds when the gene should remain inactive (2), we consider this as crosstalk. For a repressor-regulated gene, crosstalk states occur when a non-cognate repressor binds when the gene should be active (3), or if the cognate repressor fails to bind when the gene should be inactive (4). We present cognate TFs by dark gray and non-cognate ones by light gray. Activators are represented by right-inclining and repressors by left-inclining ellipses. Crosstalk states are marked by red crosses.

We define ‘crosstalk’, which potentially leads to an undesired regulatory outcome, as cases in which a binding site that should be bound by a specific TF is instead bound by a non-cognate one or remains unbound (*x*_bound_), or in which a binding site that should have been unbound is occupied (*x*_unbound_) - see Fig 1C. To quantitate the probability of these events, we use the thermodynamic model of gene regulation [22, 23, 19, 20]. A mathematical model for crosstalk for the special case in which all TFs are activators (*p =* 0) was derived and analyzed in a previous work [21]. Here, we analyze a more general model with a combination of activators and repressors. The reader can find the details of both models in the SI of this paper.

Both activity and inactivity of genes can be attained by means of either activator or repressor regulation. Accordingly, our model distinguishes between four sets of genes (see Table below and Fig 1B):

| Activity | Regulated by | Proportion of genes using this regulatory strategy |
| --- | --- | --- |
| active | activator | $a$, where $a \leq q, p$ |
| active | repressor | $q - a$ |
| inactive | activator | $p - a$ |
| inactive | repressor | $(1 - p) - q + a$ |

The probability that a particular gene *i* is in the *x*_bound_ or *x*_unbound_ crosstalk states, depends on the copy number of competing non-cognate TFs, *C*_*j*_, *j* ≠ *i* and on the number of mismatches, *d*_*ij*_, between each competing TF *j* and the regulatory binding site of gene *i*, where we assume equal energetic contributions of all positions in the binding site. Consequently, the similarity between binding sites regulated by distinct TFs is a major determinant of crosstalk. We introduce an average measure of similarity between binding site *i* and all other binding sites *j* ≠ *i* [21]:

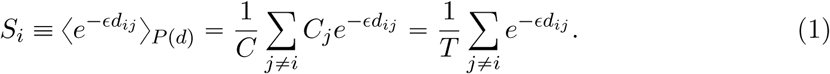

As only a subset of the genes is regulated, the summation of only the corresponding subset of TFs available to bind is taken. *S*_*i*_ is defined as the average of the Boltzmann factors, 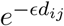, taken over the distribution of mismatch values *P* (*d*) between binding sites *I* and *j*, ∀*j*. In the last equality in Eq. (1), we assume that all available TFs are found in equal concentrations *C*_*j*_ *= C*/*T*, ∀*j*, where *C* is the total TF concentration and *T* is the number of distinct TF species available. We found that allowing different concentrations for activators and repressors does not reduce crosstalk below this equal concentration scheme (SI). We also assume full symmetry between binding sites *i*, such that *S*_*i*_ = *S* ∀*i*. A numerical analysis of a more general case with non-uniform *S*_*i*_ values can be found in SI (Fig S2). The value of *S* can be either estimated using binding site data (see below) or analytically calculated under different assumptions on the pairwise mismatch distribution *P* (*d*). In the following, we use rescaled variables: *s* = *S* · *M* for rescaled similarity between binding sites, the fraction of available TFs (*t = T* /*M*) and the rescaled total TF concentration (*c* = *C*/*M*).

We distinguish crosstalk states of genes whose desired state of activity requires unoccupied binding sites (*x*_unbound_), and those requiring occupation by a cognate regulator (*x*_bound_). *x*_unbound_ crosstalk includes the cases of an activator-regulated gene that should remain inactive as well as that of a repressor-regulated gene that should be active, both requiring an unoccupied binding site. For these genes, the cognate TF is not available to bind and any binding event by another (non-cognate) regulator is considered crosstalk. *x*_bound_ crosstalk includes both an activator-regulated gene that should be active and a repressor-regulated one that should be inactive. For these, crosstalk states occur either if the binding site remains unbound or if it is occupied by a non-cognate regulator, in which case, the regulatory state is not guaranteed. For illustration of all possible crosstalk states, see Fig 1C. Using equilibrium statistical mechanics, these crosstalk probabilities for a single gene *i* are [23, 19, 21]:

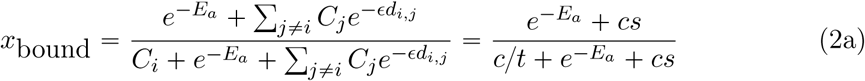

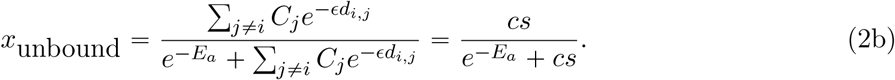

*E*_*a*_ is the energy difference between cognate bound and unbound states. The expression 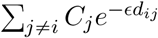 captures the sum of all interactions of binding site *i* with foreign regulators.

### Global crosstalk depends on the use of regulators

We define the global crosstalk, *X*, of a cell as the average fraction of genes found in any of the crosstalk states. For a given value of *a*, we average over different choices of *a* active genes out of the *p* activator-regulated and over different choices of *q* − *a* out of the (1 − *p*) repressor-regulated proportions. The weighted sum over these four types of contributions provides the average total crosstalk, *X*, of the whole system:

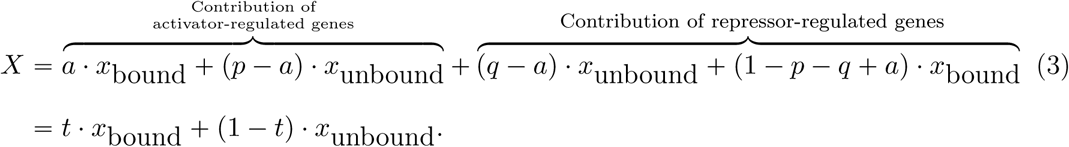

As Eq. (3) shows, *X* simply depends on the fraction of available TF species *t =* 1−*p*−*q* + 2*a*, where *t = T/M*, regardless of their role as activators or repressors. Importantly, global crosstalk does not directly depend on the fraction of active genes *q*. This is a generalization of the result obtained in [21], where the special cases of *t* = *q* (all TFs are activators) and *t* = 1 − *q* (all TFs are repressors) were studied. To obtain a lower bound on crosstalk values for given similarity, *s*, and fraction of available TFs, *t*, we substitute the expressions for *x*_bound_ and *x*_unbound_ (Eq. (2)) into Eq. (3). We then minimize *X* with respect to the total TF concentration, *c*. Such minimization is possible because global crosstalk balances between some binding sites that should be bound and others that should be unbound. For the former, higher *c* increases their chance to be bound by their cognate TFs and thus reduces crosstalk. For the latter, their cognate TF is absent and thus higher *c* increases their chance to be bound by foreign TFs, namely increases crosstalk. We then obtain the expression for minimal crosstalk:

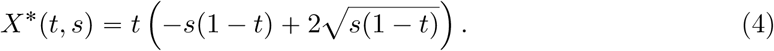

Hence, the lower bound on crosstalk *X*\* only depends on two macroscopic variables: *s* (similarity between binding sites) and *t* (fraction of available TFs). The higher the similarity *s*, the larger the resulting crosstalk *X*\*, where to first order, 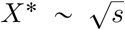 (Fig 2A). The dependence on *t* is more complicated and non-monotonic: for low *t* values, *t* < *t*\*(*s*) (we show in the SI that *t*\*(*s*) ≤ 2/3), *X*\* increases with *t*. Intuitively, the number of available TF species positively correlates with the number of crosstalk opportunities. Contrary to this intuition, for high TF usage beyond the threshold value *t*\*, we find the opposite trend, where *X*\* *decreases* with increasing TF usage, *t*. This non-monotonic dependence of *X*\* on *t* comes about because the optimal concentration *c*\*(*s, t*) is tailored specifically for each *t* value, because the relative weight of binding sites that should be bound vs. those that should be unbound, shifts with *t*. High TF usage though always comes at the cost of an exponential increase in the optimal TF concentration, *c*\*, (Eq. S4), where for high *s* values, *c*\* diverges to infinity *c*\* → ∞ (see Fig 2B). We discuss below the biological relevance of the high *t* regime.

**Figure 2:**
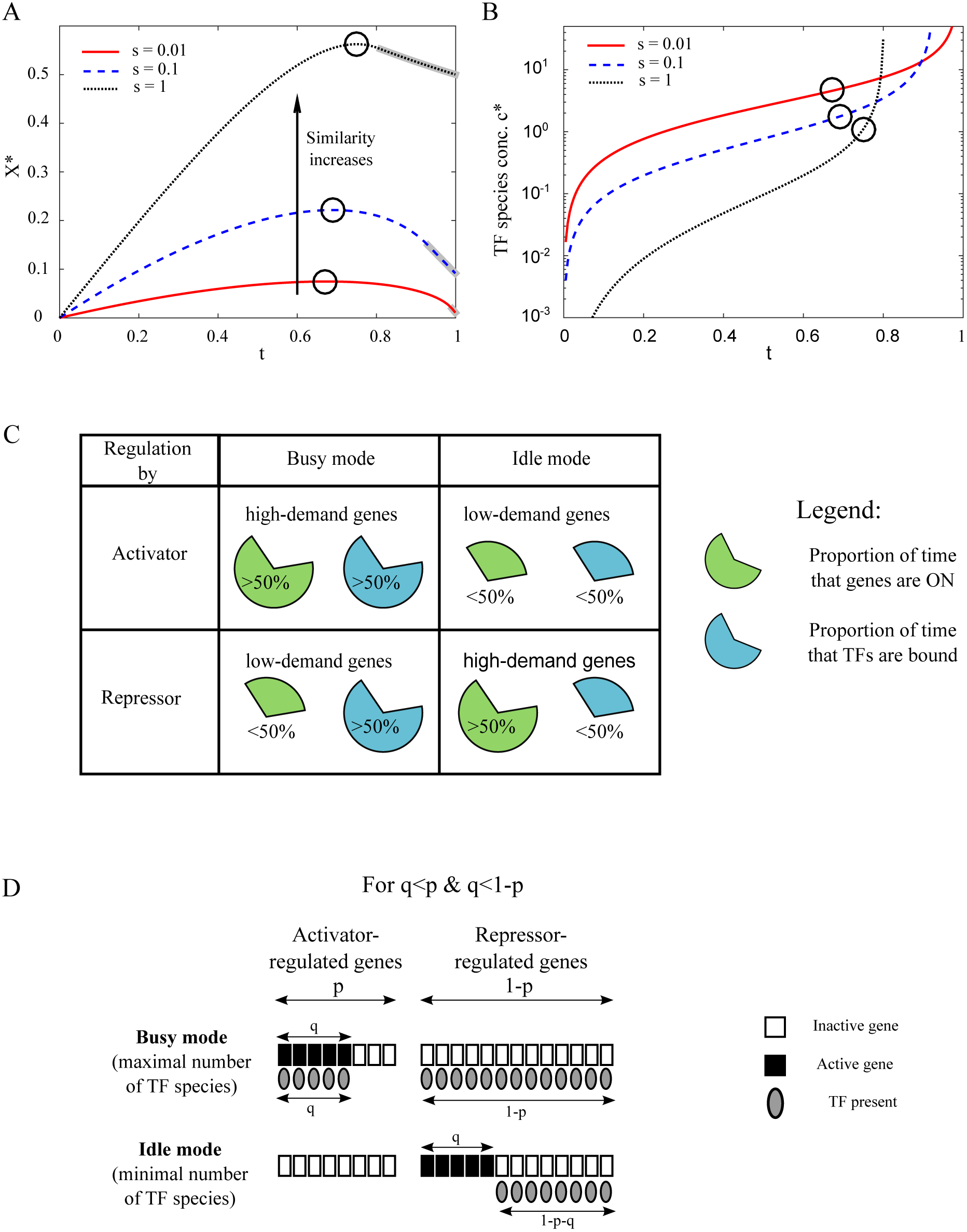
Crosstalk depends on the fraction of TFs in use, which varies between regulatory designs. **(A)** We illustrate minimal crosstalk, *X*\*, vs. *t*, the fraction of TFs in use, for different values of similarity, *s*. In most of the parameter regime (for *t* < *t*\*, *t*\* ≥ 2/3), minimal crosstalk, *X*\*, increases with *t*. Black circles denote the maxima of the curves. Crosstalk monotonically increases with similarity between binding sites. The anomalous regime where TF concentration needed to minimize crosstalk mathematically diverges to infinity, is grey-shaded around the curves. **(B)** The optimal TF concentration, *c*\*, needed to minimize crosstalk increases sharply with *t. c*\* diverges to infinity at the boundary with the anomalous regime, which for high similarity *s*, occurs already at lower TF usage *t*. Circles represent the maximal *X*\* values for each curve (as in (A)). **(C)** Different genes are expressed to different extents, where here, we grossly classify them as either high-(more than half of the time) or low-demand (less than half). If a high-demand gene is regulated by an activator or if a low-demand gene is regulated by a repressor, demand for the regulator will be high (‘busy design’). Conversely, if the same high-demand gene is regulated by a repressor and the low-demand gene is regulated by an activator, the regulator is only required for a small fraction of the time (‘idle design’). **(D)** Each of the *q* active genes and 1 − *q* inactive genes can be assigned either positive or negative regulation. We illustrate the two extremes maximizing (minimizing) TF usage: in the ‘busy’ (‘idle’) design, as many active genes as possible are assigned positive (negative) regulation and as many inactive genes as possible are assigned negative (positive) regulation. The scheme is shown for an example with the given relationship between the proportion of active genes *q*, the proportion of activator-regulated *p* and the proportion of repressor-regulated (1 − *p*); that is, *q* ≤ *p*, (1 − *p*). Other combinations are shown in SI Fig S5.

### Mode of regulation affects global crosstalk because it affects TF usage

A particular gene activity pattern can be obtained by different combinations of positive and negative regulation, yielding seemingly identical gene functionality. One may then ask whether these various TF-gene associations differ in the resulting global crosstalk. Following Eq. (4), crosstalk only depends on the fraction of available TF species, *t*, regardless of the underlying association of a gene with either activator or repressor. It is thus sufficient to consider how different regulatory strategies affect TF usage, rather than analyzing the whole network architecture, thereby significantly simplifying the analysis. Using our model, we calculate the global crosstalk for any combination of the fraction of active genes, *q*, with any mixture of activators and repressors defined by *p*, thereby covering all possible gene-regulator associations with either activators or repressors. While each point represents a fixed fraction of active genes, this model can also be used to study a varying number of active genes, by taking a distribution of points over the *q*-axis (see SI for an example). Specifically, we focus on the two extreme gene-regulator associations, which we call the ‘busy’ and ‘idle’ network designs. The ‘busy’ design means that gene regulation is operative most of the time. It is implied by the “Savageau demand rule” [2], because the gene’s default state of activity is not its commonly needed state. Under the opposite ‘idle’ design, the default state of each gene is its more commonly needed regulatory state. Hence, regulation is inoperative most of the time (see Fig 2C). Hybrids of these two extreme designs are also possible.

To represent the ‘busy’ design, we associate as much of the *q* active proportion as possible with activators, and only if the total fraction of activators is smaller than the fraction of active genes (*p* < *q*), the remaining *q* − *p* proportion is regulated by repressors. Thus the fraction of activator-regulated active genes is *a =* min (*p, q*). Conversely, under the ‘idle’ design, we associate as much of the *q* active proportion as possible with repressors. Only if the fraction of repressors is smaller than the proportion of active genes (1 − *p* < *q*), the remaining active genes pursue positive regulation, hence *a* = *q* − min ((1 − *p*), *q*). The corresponding fractions of TFs in use (including both activators and repressors) in these two extremes are then:

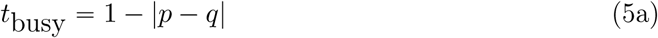

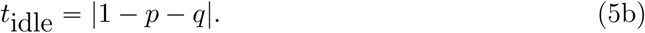

In Fig 2D, we illustrate regulation following these two extreme designs. We show that the TF assignments defined in Eq. (5) are the two extremes in TF usage. Namely, for any general regulatory scheme, the fraction of TFs needed to regulate a given fraction of genes *q* is *t*_idle_ ≤ *t* ≤ *t*_busy_ (see SI for formal proof).

In Fig 3A, we illustrate the difference in the fraction of available TFs between the two extreme designs Δ*t* = *t*_busy_ − *t*_idle =_ 1 − |*p* − *q*| − |1 − *p* − *q*| > 0, demonstrating that the ‘busy’ design always requires more regulators than the ‘idle’ design (see SI).

**Figure 3:**
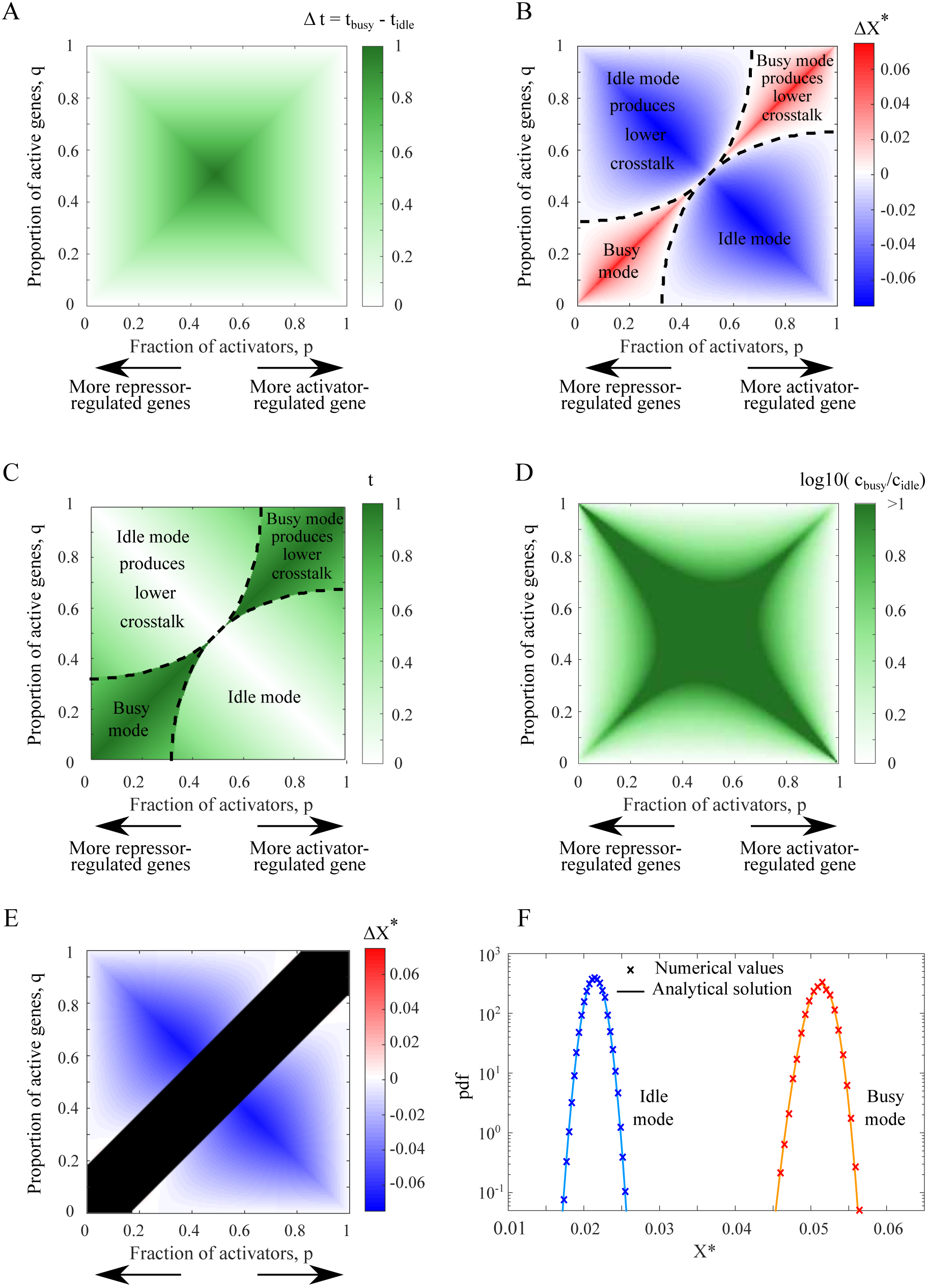
‘Idle’ design yields lower crosstalk than the ‘busy’ in a large part of the parameter regime. **(A)** The ‘busy’ design always requires more TFs compared to the ‘idle’ design. Here we illustrate Δ*t* (shown in color scale), i.e., the difference in the fraction of TFs in use between the two designs for different values of *p* and *q*. **(B)** The difference in total crosstalk 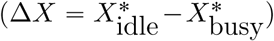 between ‘idle’ and ‘busy’ designs, shown in color scale, as a function of the fraction of activator-regulated genes *p* and relative number of active genes *q*. In a large part of the parameter regime (colored blue), lower crosstalk is achieved by following the ‘idle’ design. The ‘busy’ design is most beneficial on the diagonal *p = q* (red region), but this requires use of all TFs and comes at the cost of an enormously high TF concentration. The ‘idle’ design is most beneficial around the anti-diagonal *q* = 1 − *p*, where regulation can proceed with no TFs at all and crosstalk is close to zero. **(C)** Fraction of TFs in use (shown in color scale) when the design providing minimal crosstalk (‘idle’ or ‘busy’ as in (b)) is used, as a function of *p* and *q*. Black dashed lines mark the borders between the regions where ‘busy’ or ‘idle’ designs provide lower crosstalk. While ‘idle’ design mostly requires a minority (<50%) of the TFs, the ‘busy’ design always necessitates a majority (>50%) of TFs to be in use. *s =* 10^−2^ was used in (B)-(C). **(D)** Ratio between TF concentrations providing minimal crosstalk in either design 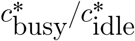. ‘Busy’ design always requires higher TF concentrations. **(E)** For higher similarity *s* between binding sites, parts of the parameter space fall into the anomalous regime where the optimal TF concentration diverges to infinity. We plot here the difference in optimal crosstalk 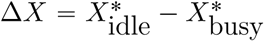 between designs for *s =* 1. Black areas denote the anomalous regime. Importantly, the region where the ‘busy’ design was beneficial for low *s* (see (B)) falls into this anomalous regime. **(F)** Analytical solution of the stochastic model for the distribution of crosstalk values, is in excellent agreement with stochastic simulation results. The distributions obtained are narrow, suggesting that their mean value is representative. Crosstalk values only depend on TF usage, regardless of the exact underlying model. Parameter values: total number of genes *M* = 3000, proportion of activator-regulated genes 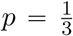, regulation probability *γ*_*i*_ = *γ =* 0.12 for ‘idle’ design and *γ*_*i*_ = *γ =* 0.92 for ‘busy’ design, with 2 · 10^6^ realizations.

Using Eq. (4), we obtain exact expressions for *X*\* under these extreme designs (see SI). In Fig 3B, we show 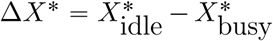, the difference in minimal crosstalk *X*\* between the two extreme designs, for all (*p, q*) combinations. We find that the ‘idle’ design yields less crosstalk in a large part of this parameter space. The ‘busy’ design still involves less crosstalk for parameter combinations centered around the diagonal *p = q*, whereas the ‘idle’ design always performs best on the anti-diagonal 1 − *p* = *q*. This is due to the fact that on the diagonal, the fraction of activators, *p*, equals exactly the fraction of genes that should be active *q*, resulting in full usage of all existing TFs, *t =* 1. On the anti-diagonal 1 − *p = q*, the fraction of genes that should be active, *q*, equals exactly the fraction of repressors 1 − *p*. Thus, the default state of all genes is the desired regulatory state requiring no TF usage at all, *t* “0, which makes the ‘idle’ design most advantageous.

In the region in which the ‘busy’ design yields the lowest crosstalk, this comes at the cost of using a larger fraction of existing TF species, as depicted in Fig 3C. The ‘idle’ design, in contrast, requires a much smaller fraction of TF species. Furthermore, the two designs differ not only in the fraction of TFs needed but also in their concentrations. To achieve the lower bound, the ‘busy’ design always requires a higher total TF concentration, *c*\* (Fig 3D).

The explanation for the alternating crosstalk advantage between the two extreme designs lies in the non-monotonic dependence of crosstalk on TF usage, *t* (Fig 2A). For *t*(*p, q*) < *t*\*(*s*), crosstalk *increases* and for *t*(*p, q*) > *t*\*(*s*), it *decreases* with *t*. Thus, for (*p, q*) combinations for which *t*_idle_ < *t*_busy_ < *t*\*, ‘idle’ design will yield lower crosstalk, whereas if *t*\* < *t*_idle_ < *t*_busy_, ‘busy’ will be more advantageous (see SI for more details). While ‘idle’ and ‘busy’ represent the two extremes, a continuum of regulatory designs interpolating between these two extremes can be defined. We show, however, that minimal crosstalk is always obtained by one of the two extremes, due to the concavity of *X*\*(*t*) (see SI).

We previously found that for some parameter combinations of similarity, *s*, and fraction of active genes, *q*, the mathematical expression for *X*\* (Eq. (4)) has no biological relevance [21]. Specifically, for similarity between binding sites which is too high 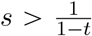, regulation is ineffective and the lower bound on crosstalk *X*\* is obtained with no regulation at all. Another biologically irrelevant regime occurs for high TF usage *t* a *t*_max_ (see SI of [21]). Then the concentration needed to obtain minimal crosstalk formally diverges to infinity *c*\* → ∞. These biologically implausible regimes put an upper bound to the total number of genes that an organism can effectively regulate [30, 21]. The results shown in Fig 3 only refer to crosstalk values obtained in the ‘regulation regime’ where *c*\* is finite and positive, 0 < *c*\* < ∞. Specifically, we find that when similarity, *s*, increases, parts of the parameter space shown in Fig 3A indeed move into the anomalous regimes. In particular, the high TF usage region around the diagonal *p* = *q*, where the ‘busy’ design outperforms in crosstalk reduction, vanishes due to this anomaly (see Fig 3E where anomalous regions are blackened). For high similarity values *s* > 5, the ‘idle’ design yields lower crosstalk in the entire biologically relevant parameter space - see SI and Fig S7.

### The distribution of crosstalk in a stochastic gene activity model

So far, we considered a deterministic model in which the numbers of active genes and available TF species were fixed, resulting in a single crosstalk value per (*p, q*) configuration. In reality, these numbers can temporally fluctuate, for example, because of the bursty nature of gene expression [31, 32]. In the deterministic model, we also assumed uniform gene usage, such that all genes are equally likely to be active. In reality, however, some genes are active more frequently than others.

To account for this, we study crosstalk in a probabilistic gene activity model. We assume independence between activities of different genes, where each gene *i, i* = 1*…M*, has demand (probability to be active) *D*_*i*_. We then numerically calculate crosstalk for a set of genes. This approach enables us to incorporate a varying number of active genes and a non-uniform gene demand and compare our results to the deterministic model studied above. To comply with its demand *D*_*i*_, each gene *i* is regulated with probability *γ*_*i*_, where *γ*_*i*_ = *D*_*i*_ if regulation is positive and *γ*_*i*_ = 1 − *D*_*i*_ if it is negative. We then obtain exact solutions for the distributions of *t* and *X*\* (Eqs. S14 in SI). In Fig 3F, we illustrate the *X*\* distributions for two values of *t*, representative of the two extreme designs. We find excellent agreement between this analytical solution and stochastic simulation results. The distribution of *X*\* is typically narrow, such that for practical purposes, the distribution mean, calculated using the deterministic activation model, serves as an excellent estimator of crosstalk values. For more details on this calculation and for approximation of the distribution width, see SI.

### Data-based crosstalk calculation

Similarity and crosstalk, considered in our analytical model, can be estimated from bioinformatic data. As direct thorough measurements of TF binding preferences are available for only a few TFs [16, 28, 29], we use statistical estimates based on multiple binding sites to which a particular TF binds (PCM) to determine its binding energetics to various sequences.

Specifically, we use data of 23 *S. cerevisiae* transcription factors collected from the scerTF database [33, 34]. PCMs are 4 × *L* matrices that provide the total number of counts for each nucleotide at each of the *L* binding site positions, taken over multiple binding sites of the particular transcription factor. They allow us to compute the mismatch energy penalties for every position and nucleotide in a given binding site sequence and then numerically calculate crosstalk.

In our theoretical model, we made several simplifying assumptions to allow for an analytical solution. In particular, we assumed uniform properties for all binding sites, assigned equal energetic contributions to all nucleotides in the sequence and assumed that all TFs regulate an equal number of genes (a single gene per TF, in the basic model). The availability of TF binding data allows us to relax these assumptions, and consider variation in binding energies and promiscuity among TFs, as well as the actual unequal energetic contributions of the different positions in each binding site.

Due to shortage of data on actual concentrations, we still assume equal concentrations for all available TFs. In addition, due to paucity of data on epistatic effects between distinct binding site positions, we still assume additivity in the energetic contributions of different positions in the sequence. The latter assumption is considered reasonable for up to 3-4 bp substitutions [16].

### Similarity values vary between genes even within the same organism

We begin by numerically calculating the similarity *s*_*i*_ between transcription factors (see Methods). In Fig 4A, we show the distribution of similarity values of genes associated with 23 *S. cerevisiae* transcription factors (top). We find a broad distribution of *s*_*i*_ values spanning over 5 orders of magnitude, where its median is around 10^−4^ − 10^−3^. This finding is in marked contrast to the full symmetry and equal *s*_*i*_ values for all TFs assumed in our analytical solution.

**Figure 4:**
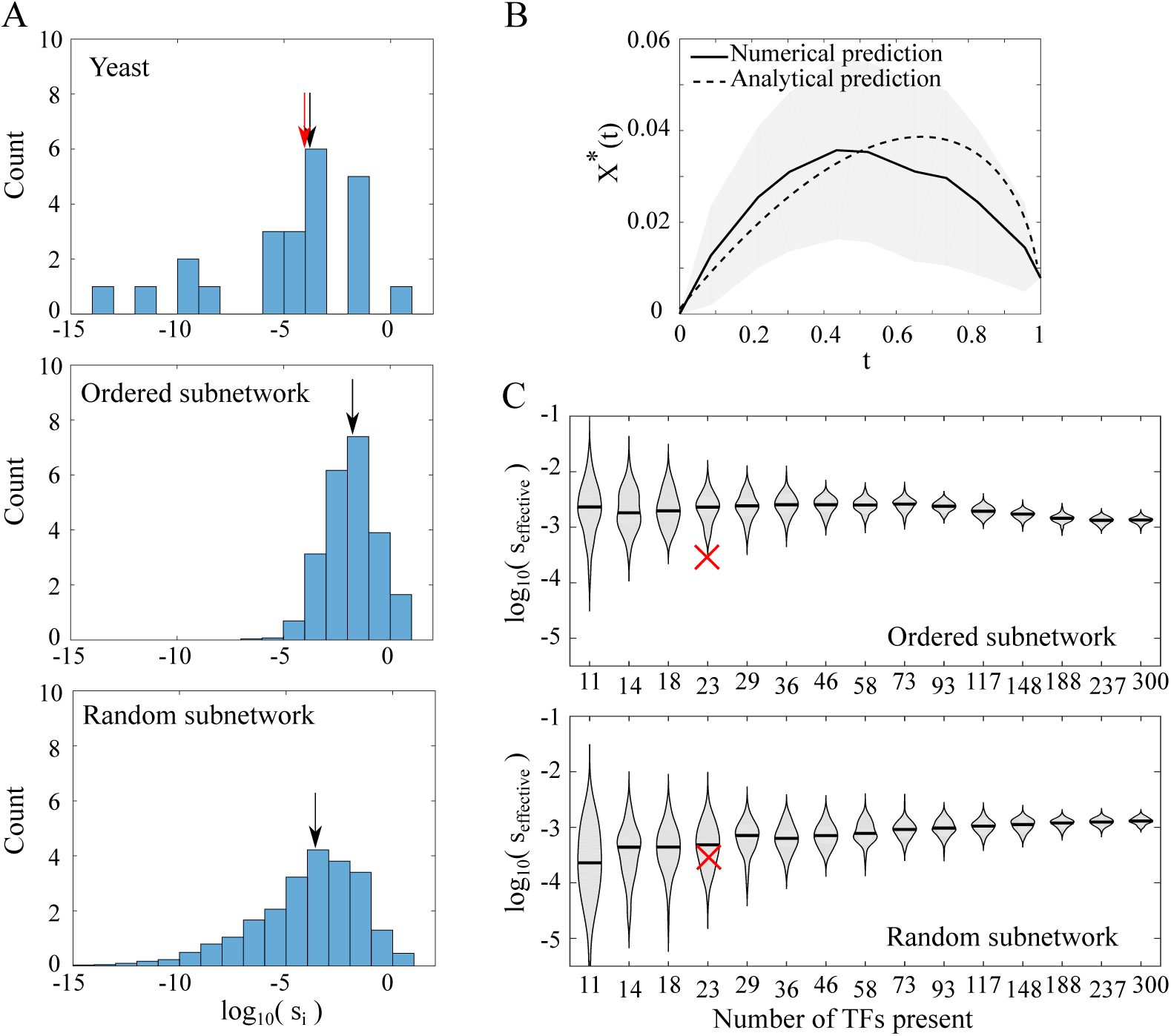
Data-based crosstalk estimates. **(A)** Inter-TF similarity values of *S. cerevisiae* TFs (top), and of synthetic data (middle and bottom) exhibit broad distributions spanning a few orders of magnitude. The distribution median values are marked by black arrows. The red arrow in the yeast data represents *s*_effective_ of the yeast data, and nearly overlaps with the distribution median. Synthetic data were created by randomly drawing PCMs representing all TFs of an artificial network. Then, sub-networks of 23 TFs were sampled by either taking the 23 most promiscuous TFs (middle) or randomly choosing them (bottom). The figures show similarity distributions amongst TFs in these artificial networks, averaged over 100 repeated draws. *s*_*i*_ values here are with respect to all TFs in the network, regardless of their (un)availability. **(B)** Numerical prediction of minimal global crosstalk depending on TF usage *t* for *S. cerevisiae* (solid line) compared to an analytical prediction based on a single *s*_effective_ value common to all genes (dashed line). This effective similarity value was chosen to provide the best fit to the numerical curve. The curves represent estimation of crosstalk for the network of all 2126 *S. cerevisiae* downstream genes regulated by the 23 TFs, for which we have PCMs. The numerical curve represents the mean over 10^3^ realizations for each *t* value, where the exact subset of available TFs was randomly drawn. The surrounding gray shadings show ±1 standard deviation around the mean. The discrepancy between numerical and analytical calculations is attributed to the broad distribution of *s*_*i*_ values for the numerical calculation, whereas the analytical calculation assumes a uniform *s*_*i*_ value for all TFs. **(C)** Violin plots of *s*_effective_ for different subnetwork sizes for *ordered* and *random* subnetworks. Ordered subnetworks are the subsets of TFs having highest similarity *s*_*i*_ with respect to the whole network. Random subnetworks include a random subset of the full network TFs. For each subnetwork, we numerically calculated crosstalk and fitted the *s*_effective_ which would best capture the crosstalk function if all TFs had a uniform *s* value. The violin plots represent distributions of effective similarity values from 100 different randomly drawn subnetworks, each coming from an independently drawn full network of 300 TFs. The red x represents the *s*_effective_ value of the 23 yeast TFs (same value as the red arrow in A). For details on the numerical calculations of similarities and crosstalk, see Methods. All violin plots exhibit broad *s*_effective_ distributions which are broadest for the smallest subnetworks, as expected. For “ordered” subnetworks, the median *s*_effective_ value is high for the small subnetworks (which were chosen to contain the most promiscuous TFs) and then slightly decreases for bigger subnetworks. For random subnetworks, the trend is opposite.

While we find that *s*_*i*_ values are very variable, the largest contributions to global crosstalk are made by the few most promiscuous TFs (those with high *s*_*i*_ values). In the following, we fit an effective similarity value that would best capture the numerically calculated crosstalk values, had all TFs had uniform *s*_*i*_ values, as in the mathematical model (denoted by red arrow in Fig 4A). In this example, we find that *s*_effective_ is almost equal to the median *s*_*i*_ value (black arrow there).

### Numerical crosstalk calculation: incorporation of a complex TF-gene interaction network

In the analytical model, we assumed that each TF regulates only a single unique gene. Yet, in real gene regulatory networks, the same TF species often regulates multiple genes and some genes are regulated by a combination of different TFs. To account for this, we expand our dataset to include all the 2126 genes [35] regulated by the 23 *S. cerevisiae* TFs for which we have PCMs and considered all possible TF-gene interactions in this set. Notably, there is high variability in the number of genes regulated by each TF. For different values of *t* (proportion of available TFs), we randomly choose a subset of TFs to be available and accordingly compute the crosstalk probabilities for all genes, accounting for all possible TF-binding site (BS) combinations. We repeat this procedure for 20 different *t* values, with 100 independent draws of available TFs for each. In the crosstalk calculation, we assume that all available TFs have equal concentrations. In contrast to the analytical calculation, where we included crosstalk contributions from all TFs, here, only binding states associated with transcription factors that are chosen to be available, are considered.

We then fit the numerically calculated crosstalk with the analytical model, where a single *s*_effective_ is used for all TFs. Fig 4B shows both the numerically calculated crosstalk and the analytically predicted one (using *s*_effective_) for this more complex interaction network (solid and dashed lines, correspondingly). The grey shading represents ±1 standard deviation around the mean value of the numerically calculated crosstalk.

### Data incompleteness could affect crosstalk estimates

Global crosstalk accounts for the combined effects of all of the organism’s TFs and binding sites. Unfortunately, data of TF binding preferences is incomplete. Moreover, the accuracy of PCMs depends on the number of known binding sites associated with the TF of interest. Due to these technical limitations, we focused on only 23 *S. cerevisiae* TFs for which >5 binding sites (per TF) are known. However, this small subset of TFs regulates one third (!) of the yeast genes. Motivated by that, we ask how representative is a crosstalk estimation of the entire network based on this small TF subset. In other words, what fraction of the TFs (or genes they regulate) would suffice to reliably estimate the global network crosstalk. This crosstalk estimation problem is further complicated by the diversity of *s*_*i*_ values we find among TFs. To generally address these questions, we simulate synthetic gene regulatory networks, each integrating 300 TFs. We simulate the binding preferences of these TFs using the PCM statistics of the 23 yeast TFs (see Methods). We then sample subnetworks of different sizes from these full networks and numerically calculate crosstalk for each subnetwork (Methods).

We sample the full networks in two manners: we either randomly choose a subset of TFs (“random subnetworks”) or deterministically select the TFs showing the highest similarity with respect to the full network (“ordered subnetworks”). The latter choice is motivated by the prior information that the few yeast TFs for which we have reliable data, are not a random subset, but rather the subset that has the largest number of binding sites. This choice is then a worst-case estimate of global crosstalk. To compare different networks on an equal basis, we estimate the effective similarity *s*_effective_ fitted for each subnetwork. Fig 4C shows the distributions and medians of *s*_effective_ values obtained, as a function of the subnetwork size. Each distribution is based on independent draws of 100 full networks. From each full network, we sample one random and one ordered subnetwork of each size.

We find, that small-size “ordered” subnetworks exhibit higher median *s*_effective_ values but narrower distributions than the “random” subnetworks, as expected. Both “ordered” and “random” subnetworks converge to the same *s*_effective_ value for the full network (of size 300). The *s*_effective_ distribution for the full size represents variation between various full networks of same size, which is significantly smaller than the variation due to limited sampling, observed for the smaller networks. As the “ordered” subnetworks deliberately include the most promiscuous TFs, their *s*_effective_ is an over-estimate of the full network measure. In contrast, we find, that *s*_effective_ estimated for random subnetworks is an under-estimate of the full network *s*_effective_. In our synthetic data, we allowed for binding site length variation among TFs (the PCM dimension). Interestingly, we find correlation between the TF’s promiscuity *s*_*i*_ and its consensus binding site length. An opposite effect is found for the length of DNA binding sites (see Fig. S11).

Considering the sufficiency of the sample size, for an “ordered” subnetwork, a sample of ∼ 50 (out of 300) TFs provides variation close to the full network measure, whereas for “random” subnetworks, a larger sample size of around ∼ 100 TFs (out of 300) is needed. Either way, we conclude that a global crosstalk estimate is possible with only a subset of the network TFs. We compare our calculated *s*_*i*_ values of yeast data (red cross) to the estimated *s*_effective_ distributions of this subnetwork size. Interestingly, the yeast estimated crosstalk value falls below the median value for both “random” and “ordered” sampling approaches. This may imply that selection to reduce crosstalk is at work, yielding similarity values which are lower than what one would expect at random [36].

## Discussion

We studied the susceptibility of different gene regulatory networks to transcriptional crosstalk. We found a lower bound on crosstalk *X*\* “*X*\*(*t, s*), which is fully determined by two macroscopic “thermodynamic-like” variables, regardless of other microscopic details of the network. These are the fractions of available TF species, *t*, and the average similarity between distinct binding site sequences, *s*. This emergent simplification enabled us to analyze crosstalk for classes of gene regulatory networks, regardless of other network details. We showed that different network designs may vary in *t*, the TF usage they require, and hence differ in the crosstalk levels they incur, even if they have the same gene activity pattern. We analyzed two extremes: a ‘busy’ design, which maximizes the use of regulators and is equivalent to the previously proposed Savageau demand rule [1] and the opposite ‘idle’ design, that minimizes the use of regulators. Interestingly, crosstalk is minimized by either of these extremes, and not by any hybrid design. We found that, in a large part of the parameter regime, crosstalk increased with *t*, and subsequently minimized by the ‘idle’ design. In the remaining part, crosstalk was minimized by the ‘busy’ design, but came at a cost of a much higher TF concentration requirement.

We also studied a stochastic gene activation variant of the model, where the number of active genes can fluctuate. We found that it is well-approximated by the deterministic activation model, because the distributions of TF availability and minimal crosstalk are typically very narrow and centered around their mean value.

Where are real organisms located in the (*t, s*) parameter space? Reports of the number of co-expressed genes greatly vary between organisms and depend on growth conditions. For example: ∼ 10,000 different genes were reported to be co-expressed in a mouse cell (<50% of total) [37, 38], 10,000-12,000 (<50%) genes were estimated to be co-expressed in human HeLa cells [39], 3300-3500 out of 4290 genes (76%-82%) were expressed in *E. coli* during exponential growth [40, 41] and 75%-80% of the genes were expressed in *S. cerevisiae* [42, 43].

Values of similarity between distinct TF binding sites, vary not only between organisms, but also between modules and distinct genes within the same organism (see Fig 4). We estimated *s*_*i*_ and the resultant minimal crosstalk values for 23 *S. cerevisiae* TFs using PCM data. We found an extremely broad distribution of single-TF *s*_*i*_ values spanning >5 orders of magnitude, with a median between 10^−4^ − 10^−3^. Global crosstalk, however, is determined by the few high-similarity TFs. To bridge the gap between the high diversity of *s*_*i*_ in real networks and our uniform *s* analytical solution, we fitted a single *s*_effective_ value which would best capture the numerically calculated network crosstalk. For the yeast data, we found that this *s*_effective_ is very close to the distribution median. Using our estimates for *s* and *t*, we estimated minimal crosstalk *X*\* for this subnetwork of *S. cerevisiae* to be in the range 0.03-0.04 (see Fig 4B), if 30%-80% of the genes are co-regulated. Our analysis showed that, for relatively low *s* values, as we found for yeast, there was a regime in the parameter space in which ‘busy’ yields the lowest crosstalk. The choice of network design that minimizes crosstalk (‘busy’/’idle’) depends on the proportion of co-regulated genes and on the proportion of activators. For organisms with high *s* values, the regime in which ‘busy’ is beneficial is actually anomalous, and hence biologically irrelevant. Such higher *s* is expected for organisms with shorter binding sites.

Binding site data is often incomplete. To assess the validity of whole-network crosstalk estimation based on a small subset of TFs, we constructed synthetic gene regulatory networks, sampled some subnetworks and then compared the *s* estimation of full and partial networks. In the *S. cerevisiae* case, we found that a full network crosstalk estimate is possible with binding information of only 16%-33% of the TFs.

Here, we used a symmetric and admittedly simplified gene regulatory network model. Our analysis determined a lower bound for crosstalk, assuming that TF concentrations are accurately tuned. In reality, TFs are not necessarily expressed and degraded at a precise time [44] and crosstalk is thus expected to be higher. Relaxation of some simplifying assumptions made in our analytical model open new research avenues for future work. Most importantly, we assumed uniform similarity values of all TFs and all BS, whereas *S. cerevisiae* data analysis showed diversity in TF properties. In principle, a distribution of *s* values can be incorporated into the model, but would significantly complicate averaging over different sets of active genes (but see a simple example in SI). Other simplifications include the averaging over gene sets of same-size as representatives of different environmental conditions, whereas, in reality, the number of expressed genes could vary between environments (e.g., growth media [40]). We averaged over all possible choices of active genes, although only some of these activity combinations occur naturally. We also assumed that every gene has a regulator, and vice versa, although this is not always the case. Hershberg and coworkers found an imbalance between genes and regulators, where orphan repressors with no genes and orphan genes with no activators, transiently exist, and could also contribute to crosstalk [45]. Relaxation of these assumptions would require a more comprehensive characterization of gene regulatory networks and co-expression patterns than is known to date.

Our study addressed a typically overlooked cost of protein production: that of regulatory interference caused by excess regulatory proteins in the dense cellular medium. This cost is distinct from the energetic burden of unnecessary protein production, which was found to delay growth [10, 11, 12, 46].

It was previously shown that transcriptional error for a single gene is minimized when its binding site is occupied [7] - a regulatory strategy equivalent to the Savageau demand rule. However, single-gene models neglected the increase in erroneous interactions that can occur following network augmentation beyond the single gene. The regulatory cost increases super-linearly with the number of molecular species and regulatory interactions and can therefore only be determined when the network is considered as whole. This would result in a different mathematical solution to minimize global crosstalk, compared to the single-gene case.

Selection to reduce global regulatory crosstalk [47, 36], was reported in previous bioinformatic studies. Our finding that similarity between *S. cerevisiae* TFs is lower than what would be obtained in a random network with similar parameters, corroborated these reports. Yet, crosstalk is not fully eliminated by selection. Despite the functional interference it causes in the short run, crosstalk is thought to promote evolvability in both gene regulatory and signaling networks in the long run [48, 49, 50, 51, 52]. However, the interplay between these two opposing effects of crosstalk, is still poorly understood.

Crosstalk reduction is one of several functional considerations shaping the evolution of gene regulatory networks. Other considerations include the network dynamical properties [53] and protein production requirements [6]. Above all, evolution is a random process and certain network designs become fixed and continue propagating [54, 55, 56, 57]. For example, new transcription factors often evolve by duplication of an existing TF followed by neo-functionalization, thereby preserving the form of regulation of the ancestral TF [58]. Taken together, a generalized model for network evolution, which would incorporate the effects of crosstalk on different time scales, alongside traditional selection on the network to achieve a certain input-output goal, remains to be formulated.

## Methods

### Distribution of *t* is approximated by a Gaussian distribution

Given that a gene *i* has its cognate TF present with a probability *γ*_*i*_ (*i* ∈(1, *M*), where *M* is the total number of genes), the distribution of available transcription factor species in the system follows Poisson-binomial distribution. This is the probability distribution of a sum of independent Bernoulli trials that are not necessarily identically distributed, each with probability *γ*_*i*_. Its mean and variance are:

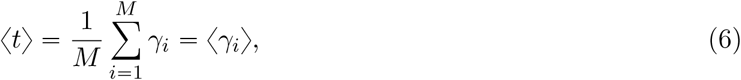

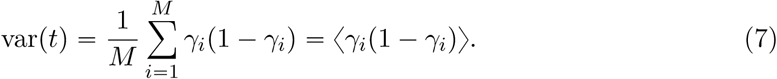

As this distribution is difficult to compute for large values of *M*, we can follow the central limit theorem and approximate it by a Gaussian distribution with the same mean and variance.

### Exact solution of probability distribution of *X*\*

For a function *X*\*(*t*), where *t* is a random variable with probability distribution *f*_*t*_(*t*), the probability distribution of *X*\*, *f*_*X*_* (*X*\*) is:

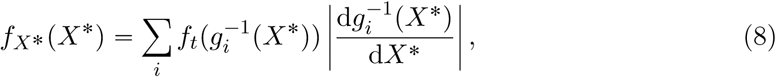

where 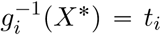 represents the inverse function of the *i*−th branch. In our case with two branches:

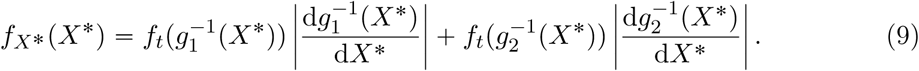

The solutions for 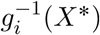 and its derivative exist for crosstalk *X*\*(*t*) and can be analytically computed. Therefore, the solution for the distribution of minimal crosstalk *f*_*X*_* (*X*\*) is also analytically known.

For regime I, the lower limit on crosstalk is *X*\*(*t*) = *t*. Its inverse is *g*^−1^(*X*\*) = *t*(*X*\*) = *X*\*, while the derivate d*g*^−1^ (*X*\*)/d*X*\* = 1. Similarly, in regime II, the lower limit of crosstalk equals *X*\*(t) = 1−*t*/(1+*αt*), the inverse function *g*^−1^(*X*\*) = *t*(*X*\*) = (1−*X*\*)/(1−*α* +*αX*\*), and its derivate d*g*^−1^(*X*\*)/d*X*\* = −(1 − *α* − *αX*\*)^−2^. The analytical solution for regime III was computed using Mathematica and the solution can be found in SI.

Using these values, one can compute *f*_*X*_* (*X*\*) for *X*\* coming from any of the three regimes.

### Stochastic semi-analytical solution of crosstalk for a random number of TFs present

For each gene *i*, we randomly draw, with probability *γ*_*i*_, that its cognate TF is available. We then obtain the proportion *t* of available TFs. As this process is stochastic, the proportion *t* differs between different realizations. Next, we compute the lower limit on crosstalk *X*\*(t) for this *t* value using the analytical solution. The crosstalk is computed in regime I, II or III, as needed. Using multiple realizations (=10^6^) of *t*, we numerically obtain the distribution of crosstalk values for values of *t* ∈ (0, 1).

### Obtaining the energy matrices from position count matrices (PCMs)

Position count matrices (PCMs) document the binding site sequences of TFs. Each element *c*_*ij*_ designates the number of known TF binding site sequences with nucleotide *i* in position We obtained the PCMs from the scerTF database for *S. cerevisiae*. Given these, we calculated the energy matrices which are needed to compute the similarity measure, in the following way: for a position *j* and nucleotide *i* ∈ {*A, C, G, T*}, we computed the energy mismatch value as 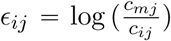, where *c*_*ij*_ is the count of nucleotide *i* in position *j* and *c*_*mj*_ = max_*i*_ *c*_*ij*_ is the nucleotide at the position *j* with the maximal count. In case of zero counts, *c*_*ij*_ = 0, where the energy *E*_*ij*_ diverges, we added a constant pseudocount *δ =* 0.1 to matrix entries.

### Some technicalities and concerns regarding PCM usage

When computing the energy matrices using PCMs, certain issues arise that could strongly bias the results if not properly addressed;

- *Inequality of total counts between positions* in PCM data. The sum of counts over all 4 nucleotides in a given PCM should be equal for all positions, but occasionally, positions with different total counts are found. As they bias our occurrence statistics (and hence our energy calculation), we used only PCMs in which the total count was equal throughout.
- *Zero counts* in the PCMs. Many PCMs include zero counts for certain nucleotides at specific positions, rendering that element of the energy matrix undefined. Here, we applied a commonly used practice of adding a pseudocount *δ* to all PCM entries. Following a previous work [21], where various *δ* values were compared to an information method (where pseudocount is not needed), we took *δ =* 0.1.
- *Count number sufficiency*. To achieve a reliable estimation of energies in the energy matrix, we only used PCMs with at least *p*_counts_ = 5 counts per position.

In total, we found 196 TF PCMs, but due to the above concerns, we considered only 23 of them in our calculations.

### Numerical computation of similarity measure using PCMs

To compute the similarity measure between binding site *k* and a transcription factor *l*, we first substituted the sequence of BS *k* by the *consensus* sequence of its cognate TF *k*. The consensus sequence is obtained by taking the most common nucleotide in each position *j*. As the given binding site and TF consensus sequence are not necessarily of the same length, we distinguished between the following cases:

- If the TF consensus sequence *l* was shorter than the binding site sequence *k*, we computed the energies for all possible overlaps of the shorter sequence with respect to the longer one. We took the minimal value to be the binding energy.
- If the TF consensus sequence *l* was longer than the binding site sequence *k*, the TF energy matrix was again slid over the binding site and energies were calculated again for every relative positioning of the two sequences. The only difference from the previous case was that energetic contributions from positions where the TF binds outside the binding site, were taken into account by averaging energies over all four nucleotides. In other words, total binding energy *E = E*_1_ + *E*_2_ is the sum of contributions from nucleotides inside (*E*_1_) and outside (*E*_2_) the binding site. The energy contribution of positions *j* outside of BS equals 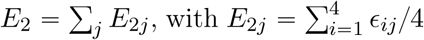 being the average binding energy at position *j* (*ϵ*_*ij*_ represent elements in the energy matrix). Here too, we computed the binding energy for all possible overlaps between the BS and TF. Then, the lowest energy was taken as the binding energy *E*_*kl*_.

This provides the matrix of binding energies *E*^*kl*^ between every binding site *k* and every TF Importantly, this binding energy is asymmetric, namely *E*^*kl*^ ≠ *E*^*lk*^. Hence, the similarity measure between binding site *k* and all other binding sites was computed as:

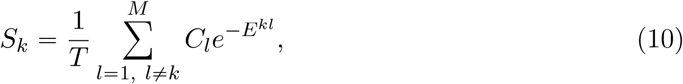

with *C*_*l*_ being the concentration of TF species *l*, and *T* the number of present TF species. In other words, the similarity *S*_*k*_ is the average Boltzmann weight, taken over all non-cognate TF binding to binding site *k*.

### Numerical computation of crosstalk given PCMs

For the numerical computation of crosstalk, we used the matrix of binding energies *E*^*kl*^ between binding site *k* and TF *l*, as per the following protocol:

1. randomly choose a subset *t* (e.g., *t =* 0.2 represents a fifth of all TFs) of all TFs that should be regulated by their cognate TF. With each realization, a different subset is chosen.
2. For gene *k*, obtain the similarity measure 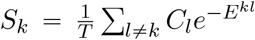, with *C*_*l*_ being the concentration of TF *l*, and *T* the number of present TF species. The concentration of absent TFs is zero. We assume that all TFs have same concentration (*C*_*l*_ = *C*), as in the analytical calculation.
3. Compute the probabilities that a crosstalk state occurs at any given gene, using the thermodynamic model. Other parameters include the energy difference between unbound and cognate state *E*_*a*_ which does not affect the final crosstalk result, and the concentration of the transcription factors, *C*.
4. Compute probability of crosstalk state occurrence for every gene and then obtain the total crosstalk *X* by summing all the individual contributions of the genes.
5. Average over a large number of realizations (we used several hundred realizations for which the average crosstalk had already converged).
6. Repeat this procedure (each with multiple realizations) using a different concentration value *C* each time. Then, pick the one that yields the lowest crosstalk value to be *X*\*(t).

### Numerical computation of crosstalk where each TF regulates multiple genes

In an actual gene regulatory network, many TFs regulate multiple genes and many genes are regulated by multiple TFs rather than the one-to-one TF-gene association we considered so far. Specifically, in our data, around 96% of the TFs regulate more than one gene. To account for that, we obtained the list of genes that are regulated by the given *S. cerevisiae* transcription factors [35].

This computation closely followed the previous procedure. The main difference was in the computation of the similarity measure of genes regulated by multiple cognate TFs. We now have multiple binding site sequences per gene (one for each cognate TF) and, as a result, more binding energies and similarity measures. To take this into account, for each gene, we averaged the similarity measures over its different binding sequences, as per the following protocol:

1. For a given gene *k*, find all the TFs that regulate it.
2. Obtain the consensus sequences of these TFs.
3. Assume each such consensus sequence represents a potential binding site sequence of gene *k* (same as in the case of only one TF regulating each gene).
4. Compute the similarity measure 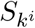 between each potential binding site sequence *i* of gene *k* and all other TFs; this is done in the same way as for one TF regulating one gene using Eq. 10.
5. Use the mean of the computed 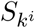 similarity measures taken over the various binding sites of gene *k* as the effective similarity of that gene.

The rest of the procedure followed the previous calculation, where each TF had only one cognate binding site: i) use the new similarity measure and compute the probability of crosstalk states, ii) do this for all genes, iii) compute the total crosstalk, iv) average over different random choices of the set of *t* genes, and v) minimize *X* with respect to the concentration *C* to obtain the *X*\*(t).

### Simulating synthetic data

To simulate synthetic data of TF binding preferences, we constructed artificial PCMs, using the statistics of the 23 yeast energy matrices, as follows. We first created the nucleotide distribution of the yeast TFs consensus sequence and then drew random realizations from this distribution to obtain a consensus sequence of each new TF. This distribution was non-uniform and was biased towards excess of A and T nucleotides. In doing so, we allowed for an unequal length of consensus sequences, as in the yeast data (using the same length distribution). We then created the distribution of the non-consensus energy values of the 23 TFs energy matrices. To construct synthetic energy matrices for these “TFs”, we drew random realizations from this distribution.

### Computing the subnetworks of synthetic data and their crosstalk

To construct a full network, we fabricated data for 300 TFs, computing the energy matrix and consensus sequence for each. We next computed the network’s matrix of binding energies 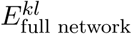 of the *l*-th TF to the *k*-th binding site, following the same procedure as for yeast data. We next formed subnetworks of this full network, by choosing a subset of TFs and taking the corresponding subset of binding energy entries, to obtain 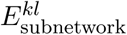. We used either randomly chosen subsets of TFs (“random networks”) or deterministically picked the subset of TFs having the highest similarity measure 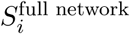 with respect to the full network. We repeated this procedure for 100 randomly drawn full networks.

We then numerically computed minimal crosstalk *X*\* for each subnetwork, following the same procedure as for yeast data.

### Comparison of numerical results to analytical expression

We fit the analytical expression for *X*\*(t) to the numerically calculated crosstalk. The main difference between the two approaches is that the analytical expression assumes uniform *s* values for all TFs, whereas the numerical approach allows for diverse *s* values. We assumed a single representative *s*_effective_ value that would best fit the numerical result. For this, we minimized the sum of squared differences over various values of *t* to find the best *s*_effective_. Distributions of *s*_effective_ values were based on 100 randomly drawn full networks from which subnetworks were sampled. For each subnetwork size, we sampled each of the full networks just once, to avoid correlations between the random subnetworks.

## Acknowledgements

We thank U. Alon and Y. Pilpel for raising the question that triggered this study. We thank Nick Barton, Yonatan Friedman, Avi Mayo, Tiago Paixão, Gašper Tkačik and Marcin Zagorski for comments on the manuscript. R. G. thanks Kirti Jain for discussions.

## Supporting Information Legends

**S1 Supporting information.**

**S2 Appendix. Maximizing crosstalk.** Mathematica notebook, showing which argument maximizes crosstalk.

**S3 Appendix. Distribution of crosstalk values.** Mathematica notebook, showing computation of distribution of crosstalk values.

