## Supporting information for "The relation between crosstalk and gene regulation form revisited"

August 5, 2019

2.2 Relation between $t$ and the choice of regulatory strategy yielding lowest crosstalk 10

**3 Maximization and minimization of TF usage: the four combinations** **11**

**4 Crosstalk expression of the two strategies** **12**

### 1 Model description

We consider a cell that has a total of $M$ transciptionally regulated genes, which can be either active or inactive. We assume that each gene is regulated by a single unique transcription factor (TF) - its cognate TF. Each gene has a short DNA binding site to which a TF can bind to affect its regulatory state. A fraction $0 \leqslant p \leqslant 1$ of the genes is regulated by activators and the remaining $(1-p)M$ genes are regulated by repressors. When no activator is bound, activator-regulated genes are inactive (or active at a low basal level) and only become active once an activator TF binds their binding site. In contrast, repressor-regulated genes are by default active, unless a repressor TF binds their binding site and inhibits their activity. We assume that different environmental conditions require the activity of different subsets of proportion $0 \leqslant q \leqslant 1$ of these genes, while the remaining fraction $1 - q$ should be inactive. As both activity and inactivity of genes can be attained by means of either activator or repressor regulation, our model distinguishes between four sets of genes: (i) $a \leqslant q, p$ activator-regulated genes which are active, (ii) $q-a$ repressor-regulated genes which are active, (iii) $p - a$ activator-regulated genes which are inactive, and (iv) $(1 - p) - q + a$ repressor-regulated genes which are inactive.

The special cases in which all the genes have the same form of regulation, either repression or activation (namely $p = 0$ or $p = 1$), were studied in a previous work [1].

We assume the system is generally at steady state, such that the required gene expression pattern does not change in time and all molecular concentrations are fixed. We then consider the average crosstalk over different gene sets of the same size. This represents a series of different gene expression patterns required in different external conditions. We assume that the system only seldom shifts from one steady state to another, such that the transient time needed for gene regulation to equilibrate following each transition is negligible. We do not consider any form of feedback exerted by the products of these genes. Rather, we assume an idealized situation in which all necessary regulators are present exactly at the time and quantity needed. Any deviation from these conditions is expected to increase crosstalk levels. Hence, our analysis refers to a lower bound of crosstalk levels.

Each gene is associated with a short regulatory DNA sequence (binding-site), to which its specialized cognate TF preferentially binds to affect its regulatory state, either positively or negatively. Although the regulatory sequences of different genes differ from each other and we assume that each TF is specific to the regulation of only one unique gene, TFs are known to have limited specificity to their DNA targets and can occasionally bind slightly different sequences, albeit with lower probability [2]. We define cases when a TF binds a non-cognate binding site or when a binding site that should have been bound remains unoccupied, as 'crosstalk', potentially leading to an undesired regulatory outcome. To quantitate the probability of these events, we use the thermodynamic model of gene regulation [3, 4, 5, 6], which asserts that the occupancy of regulatory binding sites by TFs determines the expression level of the genes associated with these binding sites. The probability of this occupancy depends on the copy number of active TF molecules available to bind and on the binding energy between the binding site and TF. This binding energy is determined by the number of mismatches between the particular binding site sequence and the consensus sequence of that TF. We assume full symmetry in the biophysical properties of the binding sites associated with different genes: all have the same sequence length and equal binding energy to their cognate TFs, and all genes have the same dynamic range of expression. Each binding site can occupy different energy levels, depending on its binding state. It is in its lowest energy level $E = 0$ if it is bound by its cognate TF. Higher energy levels are obtained if it is bound by a non-cognate TF, such that there is a mismatch between the consensus sequence of the TF and the DNA sequence of the binding site. We assume additive and equal energetic contributions of size $\epsilon$ to all nucleotides in the binding site, such that the binding energy of a TF to a sequence which differs in $d$ positions from the consensus sequence equals $\epsilon \cdot d$. Under constant external conditions, only a subset of TFs (activators and repressors) are available to bind. These TFs are needed to maintain the activity of the $q$ proportion that should be active and, simultaneously, the inactivity of the remaining $1 - q$. Unavailability means either that the TF molecules are physically absent from the cell at that time, because they were degraded, or that they are present in an inactive state and only become active in response to an external signal (e.g., via phosphorylation or other modifications). The probability that a particular gene $i$ is in either of the crosstalk states depends on the copy number of competing non-cognate TFs, $C_j$, $j \neq i$ and on the number of mismatches, $d_{ij}$ between each competing TF $j$ and the regulatory binding site of gene $i$. We distinguish crosstalk states of genes whose desired state of activity requires that their binding site remains unoccupied and those for which it should be occupied by a cognate regulator. The binding site of an activator-regulated gene that should remain inactive as well as that of a repressor-regulated gene that should be active, must all remain unoccupied. For these genes, the cognate TF is not available to bind and any binding event by another (non-cognate) regulator is considered crosstalk. For genes whose binding sites should be occupied by their cognate regulator (an activator-regulated gene that should be active and a repressor-regulated gene that should be inactive), crosstalk states occur either if the binding site remains unbound or if it is occupied by a non-cognate regulator, in which case, the regulatory state is not guaranteed. Using equilibrium statistical mechanics, the crosstalk probabilities for a single gene $i$ are [4, 5, 1]:

$$x_{\text{bound}} = \frac{e^{-E_a} + \sum_{j\neq i} C_j e^{-\epsilon d_{i,j}}}{C_i + e^{-E_a} + \sum_{j\neq i} C_j e^{-\epsilon d_{i,j}}} \tag{S1a}$$

$$x_{\text{unbound}} = \frac{\sum_{j\neq i} C_j e^{-\epsilon d_{i,j}}}{e^{-E_a} + \sum_{j\neq i} C_j e^{-\epsilon d_{i,j}}} \tag{S1b}$$

$x_{\text{bound}}$ refers to crosstalk when the binding site should be bound (by either activator or repressor) but is either unbound or bound by a non-cognate molecule. $x_{\text{unbound}}$ refers to crosstalk when the binding site should remain unbound and no cognate binder is available, but is still bound by some non-cognate molecule. $E_a$ is the energy difference between cognate bound and unbound states. The expression $\sum_{j\neq i} C_j e^{-\epsilon d_{ij}}$ then captures the sum of all interactions with foreign regulators that binding site $i$ might receive.

To account for the sum of all non-cognate interactions received by a particular binding site in a model of multiple TF species, we define an average measure of similarity between binding site $i$ and all other binding sites $j \neq i$ [1]:

$$S_i \equiv \langle e^{-\epsilon d_{ij}} \rangle_{P(d)} = \frac{1}{T} \sum_{j\neq i} e^{-\epsilon d_{ij}} = \frac{1}{C} \sum_{j\neq i} C_j e^{-\epsilon d_{ij}}. \tag{S2}$$

$S_i$ is defined as the average of the Boltzmann factors taken over the distribution of mismatch values $P(d)$ between binding sites $i$ and $j$, $\forall j$. In the last equality in Eq. (S2), we assume that all available TFs are found in equal concentrations $C_j = C/T, \forall j$, where $C$ is the total TF concentration and $T$ is the total number of available TF species. We assume full symmetry between binding sites $i$, such that each binding site $i$ has the same distribution of mismatches $d_{ij}$ with respect to all the other genes, hence $S_i = S\ \forall i$. The value of $S$ can either be estimated using binding site data (see example in Fig 4) or analytically calculated under different assumptions on the pairwise mismatch distribution $P(d)$. Following our symmetry assumptions, the crosstalk probabilities in Eq. (S1) are independent of the gene identity $i$, such that we only need to distinguish between the four different regulatory states. In the following, we use rescaled similarity defined as $s = S \cdot M$, which represents a sum of all non-cognate interactions at a binding site.

##### 1.1 For which TF usage $t^*$ is crosstalk maximized?

For a fixed $s$ $X^*(t, s)$ has a maximum at a certain $t$ value, which we denote $t^*$ (marked with a black circle on Fig 2A). We find that $t^* = t^*(s)$ and its value monotonically increases with $s$. For low similarity values ($s \to 0$), it asymptotically approaches the value of 2/3, with the limit

$$\lim_{s \to 0} t^* = 2/3. \tag{S3}$$

For $s > 0$, $t^* > 2/3$ and approaches $t \to 1$ for high $s$. See Fig S1 for illustration. Find file *argumentWhichMaximizesCrosstalk.nb* for more formal formulation.

##### 1.2 Optimal TF concentrations in the two strategies

The optimal concentration which minimizes crosstalk is $c^* = 0$ in regime I, $c^* = \infty$ in regime II, and

$$c^* = \frac{C^*}{M} = \frac{te^{-E_a}\left(s\left(st - t(st+2)\right) - \sqrt{s(1-t)}\right)}{s\left(-(st+1)^2 + st^2(st+3) + t\right)} \tag{S4}$$

in regime III. The concentration in each strategy is obtained by choosing the corresponding value of $t$, i.e., $t_{\text{busy}} = (1-p) + 2\min(p, q) - q$ and $t_{\text{idle}} = (1-p) - 2\min(1-p, q) + q$.

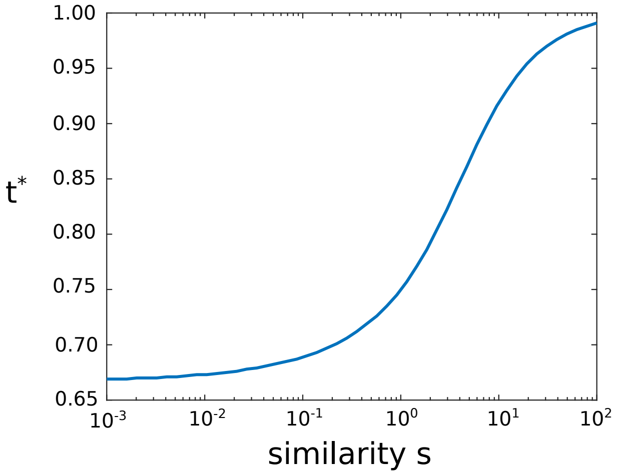

Figure S1: $t^*$ - the fraction of TFs used which maximizes crosstalk $X^*$, increases with similarity $s$. For $s \to 0$, it asymptotically approaches the value of 2/3. For large $s$, it approaches 1, such that $X^*(t)$ is an increasing function of $t$ for all but very high $t$ values.

#### 1.3 Relaxation of basic model assumptions

##### 1.3.1 Unequal TF concentrations

So far, our model assumed equal concentrations for all present TFs. What happens if we introduce different concentrations for activators and repressors? We expanded the model to allow for two concentrations, one for activators and one for repressors, while keeping the total concentrations constant. This means:

$$A \cdot C_1 + (T - A) \cdot C_1 = C, \tag{S5}$$

where $C_1$ and $C_2$ are the per species concentration of activators/repressors, $A$ and $(T - A)$ the number of activator/repressor regulated genes, and $C$ the total concentration of TFs as used in the main calculation of our model. We numerically tested which combination of concentrations $C_1$ and $C_2$ leads to lowest $X^*$ value. Surprisingly, we found that minimal crosstalk is achieved by equal concentrations for all transcription factors, activators and repressors, i.e., $C_1 = C_2$. In other words, adding an additional degree of freedom of second concentration can only increase the crosstalk values.

The intuitive explanation is that it only matters if a gene is regulated or not, but not which type of regulation it employs. This is in similarity to our previous result, where minimal crosstalk only depends on the number of available TFs, such that regulation by activators crosstalk is a mirror image of regulation by repressors alone [1].

##### 1.3.2 Non-uniform similarity values

In the basic model, we assumed uniform similarity values $s_i = s$ for all genes, which allowed us to obtain analytical solutions for crosstalk. As data show (see Fig 4), similarity values vary between genes even within the same organism. Here, we relax this simplifying assumption to test its significance. We analyze a special case with two subsets of genes, each with a different similarity value. The two subsets are of relative size $r_1$ and $r_2$ ($r_1 + r_2 = 1$), and similarity values $s_1$ and $s_2$, correspondingly. In each subset, there is a weighted proportion of regulated genes, $t_i = r_i t$ for $i \in \{1, 2\}$. We use fixed values for $s_{1,2}$ and then calculate $s = r_1 s_1 + r_2 s_2$ for each $(r_1, r_2)$ combination. As before, the total crosstalk $X^*$ is computed by summing crosstalk contributions of all individual genes and then numerically minimizing $X^*$ with respect to the TF concentration. We still allow only equal concentrations for all available TFs. In Fig S2, we plot $X^*$ vs. the proportion $r_1$ for different fractions of available TFs, $t$. We compare $X^*$ values obtained for uniform and non-uniform $s$. We find that non-uniform $s$ provides lower crosstalk than uniform $s$. This is obtained, however, at the cost of higher TF concentration $C^*$ needed for the non-uniform similarity.

### 2 Achievement of minimal crosstalk

#### 2.1 Minimal crosstalk is always obtained by one of the two extreme regulation strategies

The 'busy' and 'idle' modes explained in the main text are the two extreme regulation strategies. Intermediate strategies, where some genes follow the first strategy and others follow the second, are possible. However, minimal crosstalk is always obtained by one of the two extremes.

We denote the proportion of TF species following 'idle' and 'busy' modes by $t_1$ and $t_2$, respectively (see Fig S3). A combination of the two modes would lead to a linear combination of the fraction of TF species, $t_{\text{mixed}} = \alpha t_1 + (1 - \alpha) t_2$, with $\alpha \in [0, 1]$. Since $X^*(t)$ is a

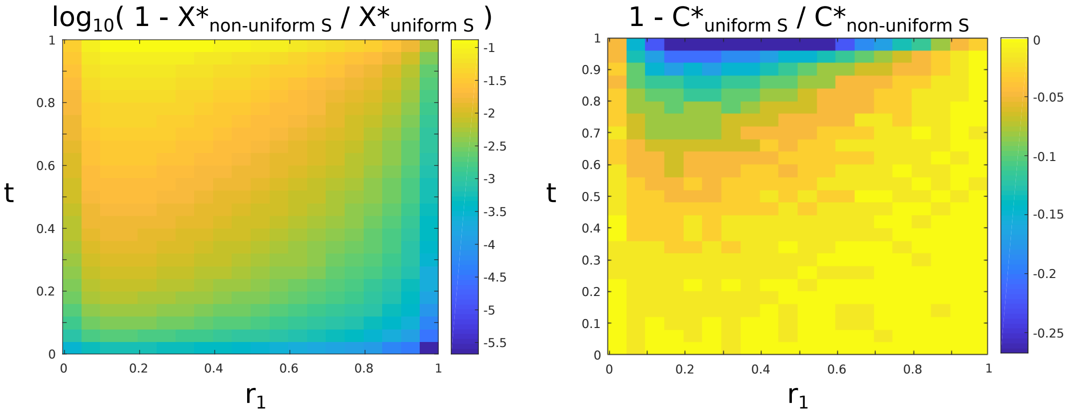

Figure S2: **Non-uniform similarity yields lower crosstalk than uniform similarity.** We plot relative change of crosstalk values (left) and concentration of TFs (right) between models of non-uniform and uniform similarities as a function of subset size $r_1$ and proportion of regulated TFs, $t$. The values of $X^*$ for non-uniform $s$ are up to 10% lower compared to uniform $s$. However, this strongly correlates with the increase in concentration. Values used: $s_1 = 5 \cdot 10^{-3}$, $s_2 = 5 \cdot 10^{-4}$.

concave function of $t$ with a single maximum, minimal $X^*$ will always be obtained at the edges of the $t$ domain, $\alpha = 1$ or $\alpha = 0$. Thus, any mixed strategy would always bring about higher crosstalk than the extreme ones $X^*(t_{\mathrm{mixed}}) \geqslant \min\left(X^*(t_1), X^*(t_2)\right)$ (see Fig S3).

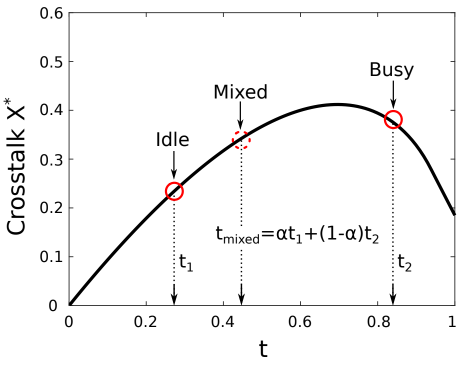

Figure S3: Due to concavity of $X^*(t)$, minimal crosstalk is always obtained by one of the two extreme modes. Crosstalk of the mixed strategy (dashed red circle) equals $X^*(t_{\mathrm{mixed}})$, where $t_{\mathrm{mixed}}$ is the linear combination $t_{1,2}$ - the proportions of TF species involved in the extreme strategies (red solid circles).

#### 2.2 Relation between $t$ and the choice of regulatory strategy yielding lowest crosstalk

The non-monotonic dependence of crosstalk on TF usage, $t$, can explain the non-trivial transition between the 'busy' and 'idle' modes in the $(p, q)$ phase space, as shown on Fig 3B. There we illustrate for each $(p, q)$ which of the two modes yields lower crosstalk. Recall that $X^*(t)$ has a maximum at $t = t^*(s)$, such that it is an increasing function of $t$ for $t < t^*$ and a decreasing function of $t$ for $t > t^*$. The relationship of $t_{\text{idle}}$ and $t_{\text{busy}}$ in regards to $t^*$ determines which strategy is more advantageous. As $t_{\text{idle}} < t_{\text{busy}}\ \forall t$ (see Eq. S12 below), it follows that if $X^*$ is increasing with $t$, idle mode is more advantageous (lower $t \rightarrow$ lower $X^*(t)$). Conversely, if $X^*$ is decreasing function of $t$, busy mode leads to lower crosstalk $X^*$ (higher $t \rightarrow$ lower $X^*(t)$). To address which mode is more advantageous, we examine $t_{\text{idle}}(p, q)$ and $t_{\text{busy}}(p, q)$ at each $(p, q)$ value (see Eq. 5 in main text). We distinguish three cases:

1. For $t_{\text{idle}}, t_{\text{busy}} < t^* \Rightarrow$ idle mode is the most advantageous strategy.

2. For $t^* < t_{\text{idle}}, t_{\text{busy}} \Rightarrow$ busy mode is the most advantageous strategy.

3. For $t_{\text{idle}} < t^* < t_{\text{busy}} \Rightarrow$ which strategy is optimal depends on exact values of $(p, q)$.

We summarize these results in Fig S4 where we show where the 3 cases lie in the phase space. The first case, where idle mode is more advantageous, occurs in the top left and bottom right corner of the phase space (white area). Conversely, $t^* < t_{\text{idle}}, t_{\text{busy}}$ holds in the bottom left and top right corner where busy mode leads to lower crosstalk (black area). The rest (gray area) belongs to the third case where it cannot be easily determined which mode is more beneficial. The boundary between idle and busy mode (red dashed line) lies entirely in the last case and can be obtained analytically by solving the equation $X^*(t_{\text{idle}}, s) = X^*(t_{\text{busy}}, s)$ for $(p, q)$. This result also intuitively explains the expansion of the region where 'busy' is advantageous when $s$ becomes smaller. Since $t^*(s)$ is a decreasing function of $s$, for smaller $s$ there is a larger $(p, q)$ region where both $t^* < t_{\text{idle}}, t_{\text{busy}}$.

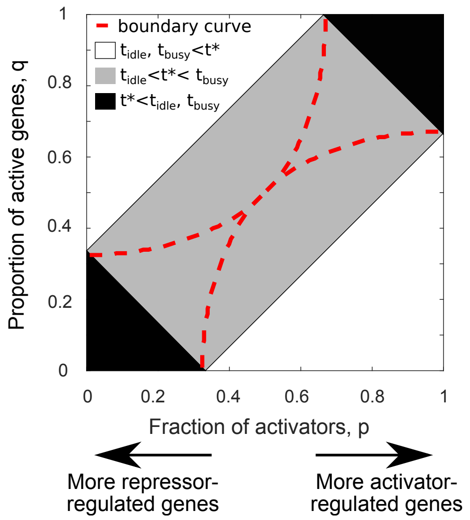

Figure S4: The three case for all combinations between $t_{\rm idle}(p,q)$, $t_{\rm busy}(p,q)$, and $t^*(s)$ in the $(p,q)$ space. Each case is shown in different color. For $t_{\rm idle}, t_{\rm busy} < t^*$ (white area), idle mode leads to lower crosstalk; for $t^* < t_{\rm idle}, t_{\rm busy}$, busy mode is more advantageous; the third region, for which $t_{\rm idle} < t^* < t_{\rm busy}$, is partitioned between the two strategies. The boundary between the optimal two modes is shown in red dashed line. For this plot we used $s = 0.01$.

### 3 Maximization and minimization of TF usage: the four combinations

The number of genes that can be associated with activators and repressors is restricted by the number of regulators of each type. When we require that a proportion $q$ of the genes is active and a proportion $p$ of the regulators are activators, we distinguish four cases depending on the relative magnitudes of these variables:

- $q < p$ and $q < 1 - p$,
- $q > p$ and $q < 1 - p$,
- $q < p$ and $q > 1 - p$,
- $q > p$ and $q > 1 - p$.

In Fig S5, we illustrate how TF usage is maximized and minimized in each of these cases (one of them appeared as Fig 2D in the main text).

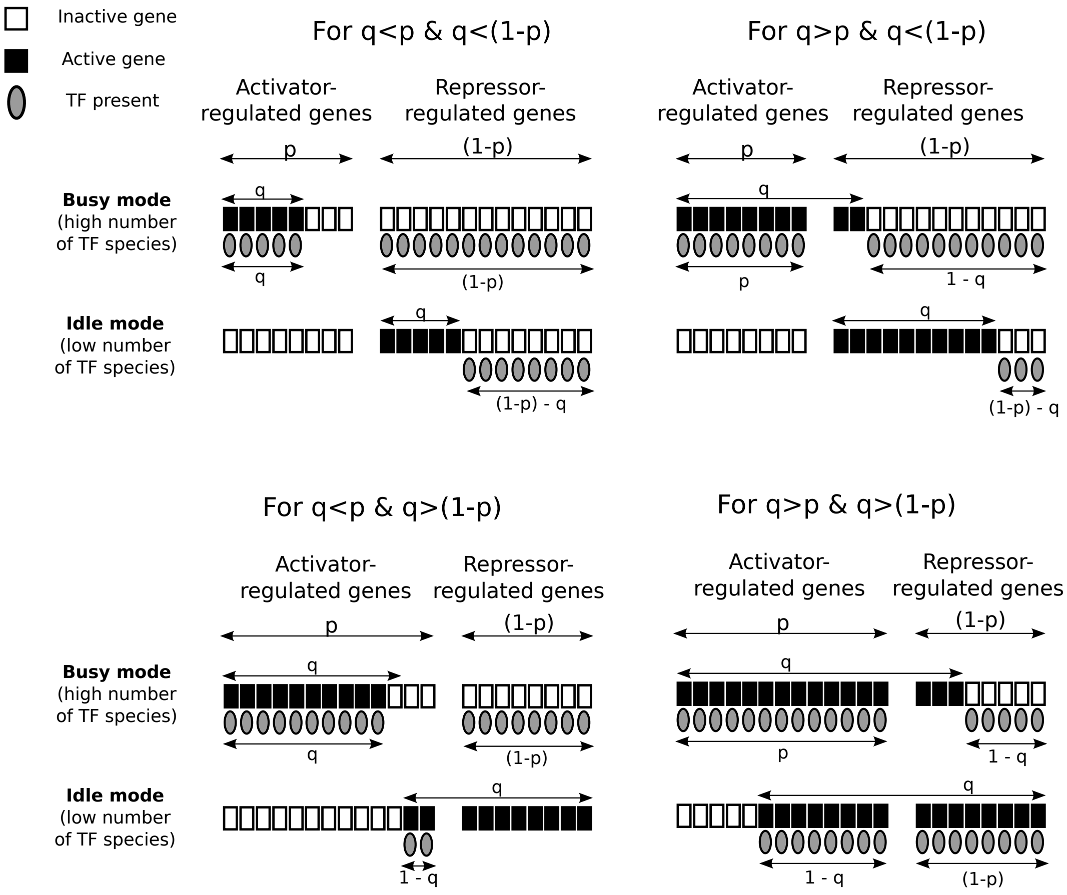

Figure S5: Minimization and maximization for four different combinations of active genes $q$ and activator regulated genes $p$.

### 4 Crosstalk expression of the two strategies

#### 4.1 Regime III

##### 4.1.1 Crosstalk expression for 'busy' strategy

In the busy mode, the proportion of active genes that are regulated by activators is $a = \min(p, q)$. The proportion of TFs involved in the busy strategy equals:

$$t_{\text{busy}} = (1 - p) + 2\min(p, q) - q. \tag{S6}$$

To obtain the lower limit on crosstalk, we use $t_{\text{busy}}$ in the equation for $X^*(t)$ and obtain:

$$X^*_{\text{busy}} = (1 - p + 2\min(p,q) - q) \cdot \left(-s(p + q - 2\min(p,q)) + 2\sqrt{s\,(p + q - 2\min(p,q))}\right) \tag{S7}$$

##### 4.1.2 Crosstalk expression of 'idle' strategy

Similarly, in the 'idle' mode, the proportion of active genes that are regulated by activators is $a = q - \min(1-p, q)$. The proportion of involved TFs is then:

$$t_{\text{idle}} = (1 - p) - 2\min(1-p, q) + q, \tag{S8}$$

and the lower bound on crosstalk in the idle mode equals:

$$X^*_{\text{idle}} = (1 - p - 2\min(1-p,q) + q) \cdot \left(-s(p - q + 2\min(1-p,q)) + 2\sqrt{s\,(p - 1 + 2\min(1-p,q))}\right). \tag{S9}$$

#### 4.2 Regimes I and II

##### 4.2.1 Crosstalk expression of both strategies in regime I

The lower limit on crosstalk in regime I is described by $X^*(t) = t$ and:

$$X^*_{\text{busy}} = (1 - p) + 2\min(p,q) - q, \tag{S10a}$$

$$X^*_{\text{idle}} = (1 - p) - 2\min(p,q) + q. \tag{S10b}$$

##### 4.2.2 Crosstalk expression of both strategies in regime II

The lower limit on crosstalk in regime II is described by $X^*(t) = 1 - t/(1 + st)$ and:

$$X^*_{\text{busy}} = 1 - \frac{(1-p) + 2\min(p,q) - q}{1 + s\left[(1-p) + 2\min(p,q) - q\right]}, \tag{S11a}$$

$$X^*_{\text{idle}} = 1 - \frac{(1-p) - 2\min(p,q) + q}{1 + s\left[(1-p) - 2\min(p,q) + q\right]}. \tag{S11b}$$

### 5 Proportion of TFs is always higher in busy mode

The proportion of TFs in busy mode is always higher than in idle mode. This can be easily shown by:

$$\begin{aligned}\Delta t &= t_{\text{busy}} - t_{\text{idle}} \\ &= ((1-p) + 2\min(p,q) - q) - ((1-p) - 2\min(1-p,q) + q) \\ &= -2q + 2\min(1-p,q) + 2\min(p,q),\end{aligned} \tag{S12}$$

We distinguish between four cases:

- $q < p$ and $q < 1-p \Rightarrow \Delta t = -2q + 2q + 2q = 2q \geqslant 0$,
- $p < q$ and $1-p < q \Rightarrow \Delta t = -2q + 2(1-p) + 2p = 2(1-q) \geqslant 0$,
- $p < q < 1-p \Rightarrow \Delta t = -2q + 2q + 2p = 2p \geqslant 0$,
- $1-p < q < p \Rightarrow \Delta t = -2q + 2(1-p) + 2q = 2(1-p) \geqslant 0$.

In all cases, the difference $\Delta t > 0$, which shows that for any value of parameters, the proportion of TFs is always larger (or equal in the extreme case of $p \in \{0,1\}$) in 'busy' mode compared to 'idle'.

#### 5.1 Idle mode minimizes, busy mode maximizes the number of transcription factor species $t$.

The busy and the idle mode maximize and minimize the fraction of TF species, respectively. This can be shown by taking a system, where a proportion of $p$ genes is activator-regulated, while the rest $(1-p)$ is repressor-regulated. Within the activator-regulated genes, $k_1$ are active and $k_2 = p - k1$ are inactive. Moreover, within $(1-p)$ repressor-regulated genes, $k_3$ are active genes and $k_4 = (1-p) - k_3$ inactive genes. Within these, several constraints exist:

- $k_1 + k_2 = p \quad \rightarrow \quad k_2 = p - k_1$,
- $k_1 + k_3 = q \quad \rightarrow \quad k_3 = q - k_1$,

- $k_3 + k_4 = 1 - p \quad \rightarrow \quad k_4 = (1-p) - k_3 = (1-p) - q + k_1$,

where $q$ is the proportion of active genes. The total proportion of TF species is $t = k_1 + k_4 = 2k_1 + (1-p) - q$. Of course, due to the definition of the system, it holds:

- $k_1, k_2 \leqslant p$,
- $k_1, k_3 \leqslant q$,
- $q - k_1 \leqslant 1 - p$.

In attempt to see how the total proportion of TF species changes if we change different parameters of the system ($k_i$) with fixed $q$ and $p$, we compute the derivative of $t$:

$$\frac{\partial t}{\partial k_1} = 2. \tag{S13}$$

Therefore, the change of TFs with increasing $k_1$ is linear and 4 different scenarios exist. For each, the constraints described above must be met. Therefore:

1. if $q < 1 - p$ and $q > p$:
   - $t$ is minimized by $k_1 \rightarrow 0 \Rightarrow k_2 = p, k_3 = q, k_4 = (1-p) - q$, which is the *idle mode*,
   - $t$ is maximized by $k_1 \rightarrow p \Rightarrow k_2 = 0, k_3 = q - p, k_4 = 1 - q$, which is the *busy mode.*
2. if $q < 1 - p$ and $q < p$:
   - $t$ is minimized by $k_1 \rightarrow 0 \Rightarrow k_2 = p, k_3 = q, k_4 = (1-p) - q$, which is the *idle mode*,
   - $t$ is maximized by $k_1 \rightarrow q \Rightarrow k_2 = p - q, k_3 = 0, k_4 = 1 - p$, which is the *busy mode.*
3. if $q > 1 - p$ and $q > p$:
   - $t$ is minimized by $k_1 \rightarrow q - (1-p) \Rightarrow k_2 = 1 - q, k_3 = 1 - p, k_4 = 0$, which is the *idle mode*,

- $t$ is maximized by $k_1 \to p \Rightarrow k_2 = 0, k_3 = q - p, k_4 = 1 - q$, which is the *busy mode.*

4. if $q > 1 - p$ and $q < p$:

   - $t$ is minimized by $k_1 \to q - (1 - p) \Rightarrow k_2 = 1 - q, k_3 = 1 - p, k_4 = 0$, which is the *idle mode*,
   - $t$ is maximized by $k_1 \to q \Rightarrow k_2 = p - q, k_3 = 0, k_4 = 1 - p$, which is the *busy mode.*

This formally proves what was graphically shown on Fig S5: minimization of TF proportion is achieved in idle mode while the maximization is obtained in busy mode.

#### 6 For sufficiently high similarity measure, idle strategy always leads to a lower crosstalk limit $X^*$

For some parameter combinations of similarity $s$ and fraction of regulated genes $t$, the mathematical result of the lower bound on crosstalk $X^*$ has no biological relevance: (i) for sufficiently high similarity measure, regulation is ineffective and the lower bound on crosstalk $X^*$ is obtained by no regulation (zero concentration of TFs, $C^* = 0$), and (ii) for high TF usage, the optimal concentration which minimizes the lower bound on crosstalk $X^*$ diverges. Therefore, when only considering the biologically relevant regime, where ($0 < C^* < \infty$), the size of area where busy mode leads to lower crosstalk limit (Fig 3B red area) decreases with increasing similarity measure (Fig 3E).

In Fig S6, we plot the difference in optimal crosstalk $\Delta X^* = X^*_{\text{idle}} - X^*_{\text{busy}}$, where black areas denote the anomalous regime. With increasing similarity, the anomalous regime grows and covers an increasingly larger portion of the phase space. For sufficiently high similarity values, the idle strategy will always lead to the most optimal crosstalk for any value of $(p, q)$. Fig S7 shows both the minimal (blue) and maximal (red) value of $\Delta X^* = X^*_{\text{idle}} - X^*_{\text{busy}}$ over the whole space of $p \in \{0, 1\}$ and $q \in \{0, 1\}$ as a function of similarity $s$. Values of $\Delta X^* < 0$ mean that the idle mode leads to a lower crosstalk limit and vice versa for $\Delta X > 0$. Therefore, when the busy mode completely vanishes and the idle mode is the one that always yields lower crosstalk ($\Delta X^* < 0$ for all $(p, q)$ values), the maximal value of $\Delta X^*$ will be negative; when $\max_{p,q} \Delta X^* = 0$, the similarity value is such that the busy mode completely vanishes. That happens at $s_{\text{vanishing}} \approx 5$, which is far above the values of real organisms – Fig 4A in the main text shows that similarity values of *S. cerevisiae* range between $s \approx 10^{-5} - 10^{0}$.

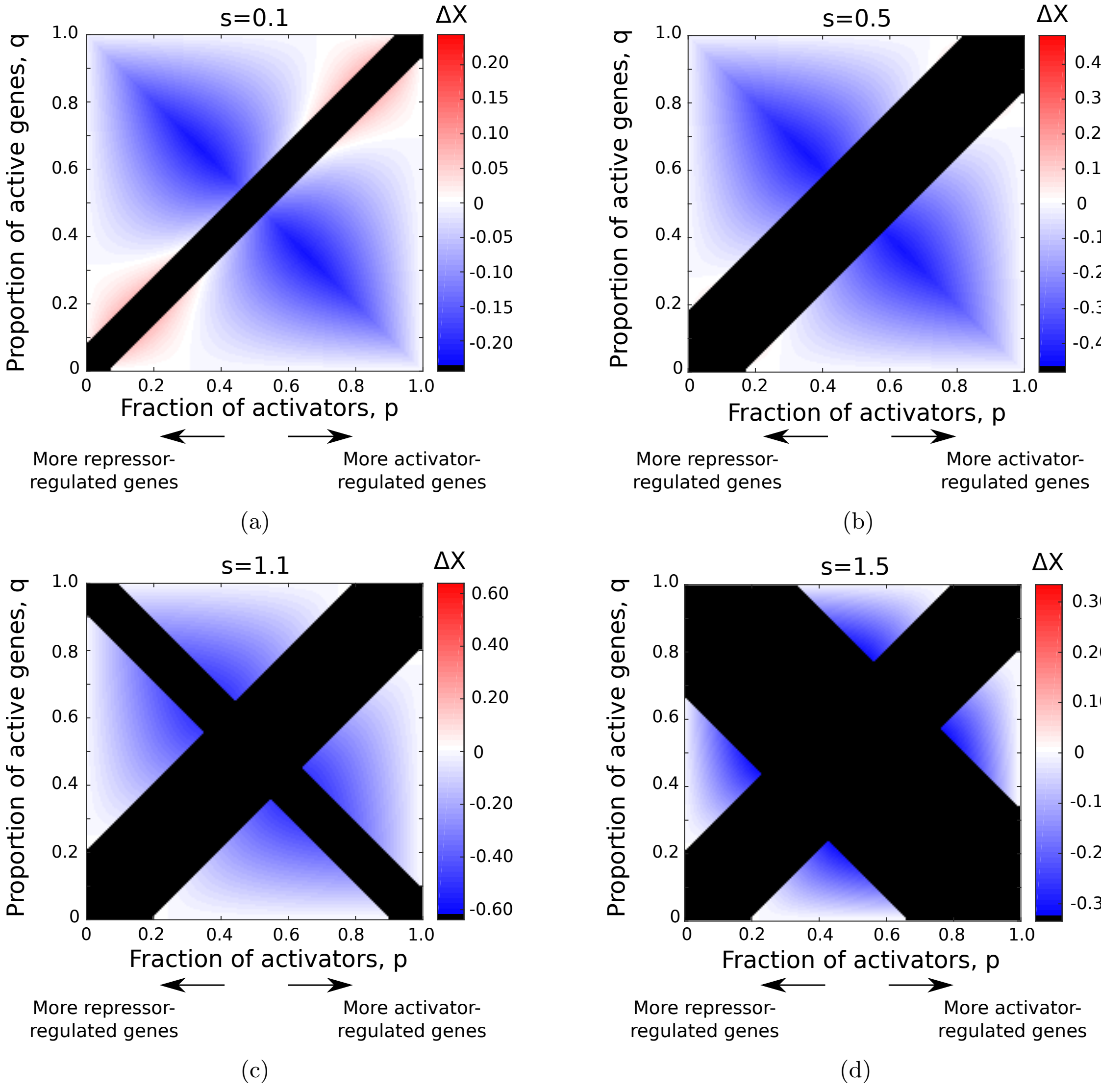

Figure S6: **The anomalous regions where crosstalk cannot be minimized, grow as the similarity $s$ increases.** Difference in optimal crosstalk $\Delta X^* = X^*_{\text{idle}} - X^*_{\text{busy}}$, where black areas denote the anomalous regime. Different values of rescaled similarity were used: (**a**) $s = 0.1$, (**b**) $s = 0.5$, (**c**) $s = 1.1$, (**d**) $s = 1.5$.

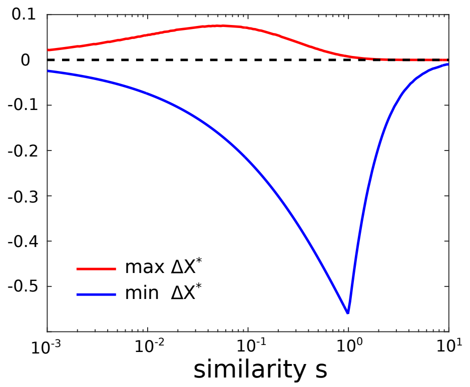

Figure S7: **The differences between regulatory strategies depend on s.** We plot the minimal (blue) and maximal (red) value of $\Delta X^* = X^*_{\text{idle}} - X^*_{\text{busy}}$ over the entire region of $p \in \{0, 1\}$ and $q \in \{0, 1\}$ as a function of similarity, $s$. The point where the maximal value (red) becomes negative (at $s \approx 5$), is where the busy mode completely vanishes and idle mode leads to lowest crosstalk (for any $(p, q)$).

### 7 Probabilistic gene activity model

#### 7.1 Probabilistic model description

So far, we considered a deterministic model in which the numbers of active genes and available TF species were fixed, resulting in a single crosstalk value per $(p, q)$ configuration. In reality, these numbers can temporally fluctuate, for example, because of the burst-like nature of gene expression [7, 8]. In the deterministic model, we also assumed uniform gene usage, such that all genes are equally likely to be active. In reality, some genes are active more frequently than others.

To account for this, we study the following crosstalk in a probabilistic gene activity model. We assume independence between activities of different genes, where each gene $i$ has demand (probability to be active) $D_i$. We then numerically calculate crosstalk for a set of genes. This approach enables us to incorporate a varying number of active genes and a non-uniform gene demand and compare our results to the deterministic model studied above.

Assume that to comply with its demand $D_i$, each gene $i$ is regulated with probability $\gamma_i$, $i = 1...M$, where $\gamma_i = D_i$ if regulation is positive and $\gamma_i = 1 - D_i$ if it is negative. We assume that the TF species needed for these genes are available in the cell. The distribution

$f_t(t)$ of the fraction of TF species follows Poisson-Binomial distribution [9] with mean and variance of:

$$\langle t \rangle = \frac{1}{M} \sum_{i=1}^{M} \gamma_i \tag{S14a}$$

$$\text{var}(t) = \frac{1}{M} \sum_{i=1}^{M} \gamma_i (1-\gamma_i). \tag{S14b}$$

Assuming that the number of genes is large, $M \gg 1$, the central limit theorem applies here: we approximate the probablity distribution of $t$, $f_t(t)$, by a Gaussian distribution with the mean value and variance as given with Poisson-Binomial distribution (mean value $\langle \gamma_i \rangle_i$ and standard deviation $\sigma_t = \langle \gamma_i(1-\gamma_i) \rangle_i$, with $\langle \cdot \rangle_i$ representing the average over all genes) [9].

Since minimal crosstalk $X^*$ is a function of $t$ (Eq. 4, main text), $X^*$ becomes a random variable and its distribution reads:

$$f_{X^*}(X^*) = \sum_l f_t(g_l^{-1}(X^*)) \left| \frac{\mathrm{d}g_l^{-1}(X^*)}{\mathrm{d}X^*} \right|, \tag{S15}$$

where $g_l^{-1}(X^*) = t_l$ represents the $l-$th branch of the inverse function (for some $X^*$ values there exist two solutions $t_l$ that satisfy the inverse equation) [9]. The solutions for $g_l^{-1}(X^*)$ and its derivative exist and can be analytically computed, which enables us to solve for the distribution of crosstalk $f_{X^*}(X^*)$:

1. Region I: $f_X^*(X^*) = f_t(X^*)$
2. Region II: $f_X^*(X^*) = f_t\left(1 - \frac{X^*}{1-\alpha+\alpha X^*}\right) \cdot (\alpha - 1 - \alpha X^*)$
3. Region III: see attached Mathematica file *DistributionOfCrosstalkRegion3.nb*,

where $f_t(t)$ is the distribution of TF usage values $t$.

As the exact calculation of the distribution $f_{X^*}(X^*)$ is difficult, we can often use the following useful approximation. If the distribution is narrow enough, such that $\sigma_T/\langle T \rangle \ll 1$, we can approximate the expected value of crosstalk by the deterministic value of crosstalk for an expected value of available TF fraction:

$$\langle X^*(t)\rangle \approx X^*(\langle t\rangle), \tag{S16}$$

where the computation of both $\langle t\rangle$ and $X^*(\langle t\rangle)$ is straightforward given $\gamma_i$. We discuss below the conditions under which this approximation holds. The distribution of $X^*$ is typically narrow, such that for practical purposes the distribution mean provides a very good estimator of crosstalk values.

#### 7.2 Approximations

In our stochastic model, a gene $i$ is regulated with probability $\gamma_i, i = 1, \ldots, M$. Above, we stated (i) that the distribution of TFs in use, $t$, can be well approximated by a Gaussian distribution. Furthermore, if (ii) in the regime where $X^*$ is linear in $t$ and $\sqrt{var(t)}/\langle t\rangle \ll 1$, one can approximate the expected value of crosstalk by the deterministic value of crosstalk for an expected value of total number of TF: $\langle X^*(t)\rangle \approx X^*(\langle t\rangle)$. The third claim is that if $X^*$ is linear in $t$ ($\partial X^*/\partial t \approx$ const.), one can also approximate well the distribution of crosstalk with a Gaussian distribution having the same mean and variance as those of the $t$ distribution, just rescaled and translated by the slope and constant factor of the linear transformation of $X^*(t)$.

The Gaussian approximation of distribution of $t$ follows from the central limit theorem. A numerical example is shown in Fig S8 (a) and (c).

The approximation (ii) uses linearity to show that $\langle X^*(t)\rangle \approx X^*(\langle t\rangle)$ holds:

$$\text{If } \ X^*(t) \approx \alpha t + \beta \quad \Rightarrow \quad \langle X^*(t)\rangle = \langle \alpha t + \beta\rangle = \alpha\langle t\rangle + \beta = X^*(\langle t\rangle). \tag{S17}$$

The linearity assumption is fulfilled for $t$ values that are much lower than $t^*$ ($t^*$ being the value at which $X^*$ reaches maximum). For visual example of linearity of $X^*(t)$ for $t < t^*$, see Fig 2A. Moreover, by using the Taylor expansion of crosstalk $X^*(t)$ for a small deviation of $t$ around its mean $\langle t\rangle$, we show that deviations around the expected value are small and often negligible. Therefore, if $\sqrt{var(t)}/\langle t\rangle \ll 1$ holds, one can look at a representative deviation from the mean value and write (assuming the solution for $X^*(t)$ is in biologically plausible regime III);

$$\begin{aligned} X^*(t) = X^*(\langle t\rangle) + \delta X^*(t) \approx X^*(\langle t\rangle) + \frac{\partial X^*(\langle t\rangle)}{\partial t}\delta t \\ = X^*(\langle t\rangle) + \left[2\langle t\rangle s - s + 2\sqrt{s(1-\langle t\rangle)} - \frac{s\langle t\rangle}{\sqrt{s(1-\langle t\rangle)}}\right]\delta t \\ = X^*(\langle t\rangle) + \underbrace{\left[\frac{X^*(\langle t\rangle)}{\langle t\rangle} - \langle t\rangle s\left(\frac{1}{\sqrt{s(1-\langle t\rangle)}} - 1\right)\right]\delta t}_{\delta X^*} \end{aligned} \tag{S18}$$

Fig S9 shows the relative error $\delta X^*(\langle t\rangle)\delta t/X^*(\langle t\rangle)$ of our approximation. We use a representative values of $M = 2500$ and $\gamma_i = 0.5$, leading to the standard deviation of $t$ being $\sqrt{var(t)} = 10^{-2}$. We take this number to also be a variation in the number of TF species present: $\delta t = 10^{-2}$. Any larger values of $M$ or any other values of $\gamma_i$ will lead to lower error. The relative error is indeed very small. The exceptions are the values close to $t = 1$, which fall out of regime III into anomalous regime II, and values close to $t_0 = 0$, which still take a small relative error of $\delta X^*(\langle t\rangle)\delta t/X^*(\langle t\rangle) = 20\%$ for $\langle t\rangle = 0.05$. Furthermore, if the third claim of $X^*(t)$ linearity with respect to $t$ holds, we can approximate the distribution of $X^*(t)$ by a Gaussian. As the distribution of $t$ is Gaussian, to a good approximation, a linear transformation of a Gaussian distribution also leads to a Gaussian distribution of $X^*(t)$. Fig S8 (a-b) shows an example of distribution of $t$ and $X^*$. There, the probabilities $\gamma_i$ give the average proportion of TF species $\langle t\rangle = 0.5$, which gives values of crosstalk that are far from the maximum of $X^*$. The assumption of linearity is justified on the example shown and Gaussian approximation for $f_{X^*}(X^*)$ gives good results. The expected values of crosstalk $\langle X^*(T)\rangle$ and the crosstalk of the expected value of TF species $X^*(\langle T\rangle)$ have a very small relative difference (in the order of 0.01%), which is the consequence of small ratio $\sqrt{var(t)}/\langle t\rangle = 2\% \ll 1$.

On the other hand, Fig S8 (c-d) shows the distribution of $t$ and $X^*$, where the probabilities $\gamma_i$ give the average proportion of TF species $\langle t\rangle = 0.66$ close to the maximum of $X^*$ around $t \approx 2/3$, where the linearity assumption does not hold. We see that Gaussian approximation for $f_{X^*}(X^*)$ is not valid anymore. Even though the linearity assumption does not hold, the expected value of crosstalk $\langle X^* \rangle$ and the crosstalk of the expected value of the relative number of TF species $X^*(\langle T \rangle)$ are again very close (relative difference of 0.03%).

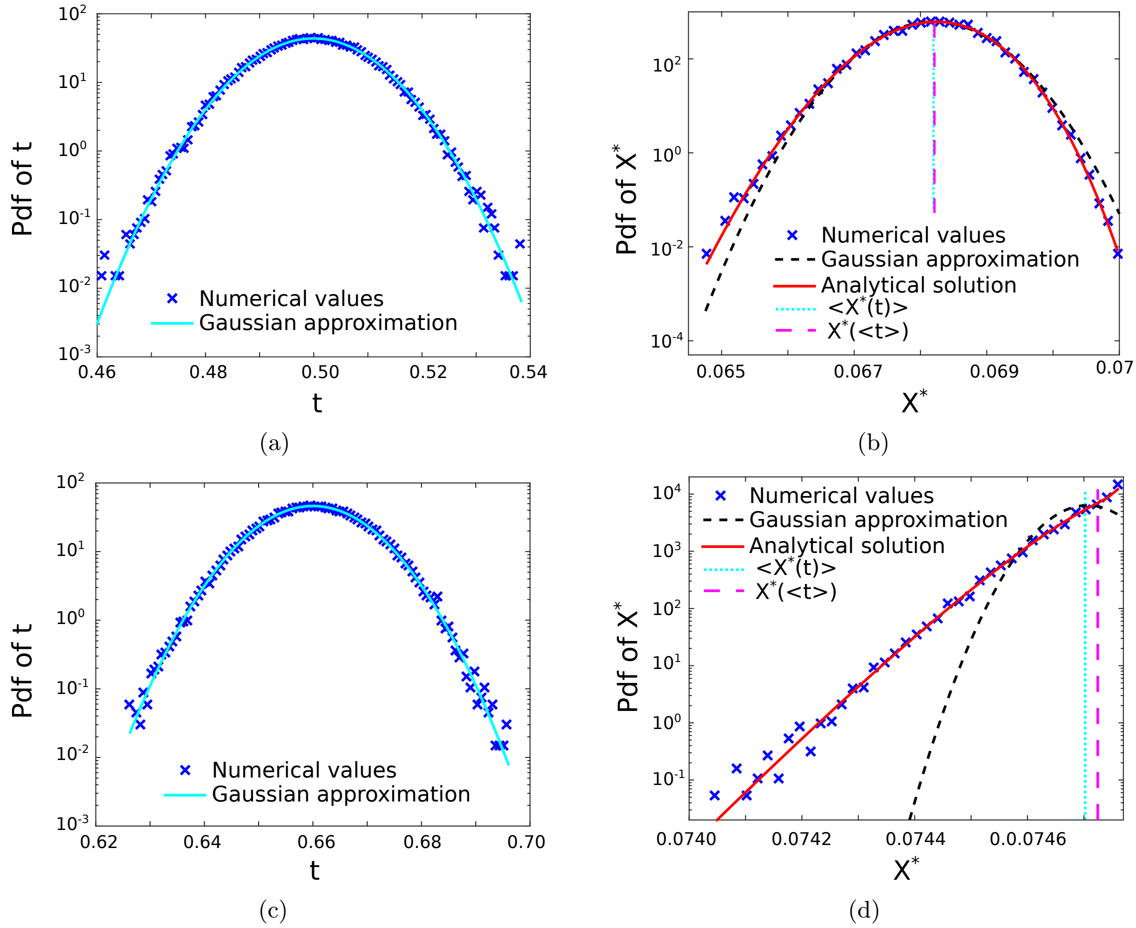

Figure S8: $t$ **distribution is always well-approximated by a Gaussian. If** $\langle t \rangle$ **is far apart from** $t^*$**,** $X^*$ **distribution is also well approximated by a Gaussian.** We plot the distributions of $t$ ((**a**), (**c**)) and $X^*(t)$ ((**b**), (**d**)) in two cases. The vertical lines in (**b**) and (**d**) (dotted and dash-dotted) represent $X^*(\langle t \rangle)$ and $\langle X^*(t) \rangle$, correspondingly. We find an excellent match between their values, even in the worst case scenario that $X^*(t)$ is far from being Gaussian (d). Small discrepancies between the analytical solution and numerical simulation is due to the finite number of iterations in the simulation. Parameter values: $s = 0.01$, $M = 3000$, $p = 1/3$, in (**a**) and (**b**) $\gamma_\mathbf{i} = \mathbf{0.66}$; in (**c**) and (**d**) $\gamma_\mathbf{i} = \mathbf{0.66}$.

##### 7.3 The probabilistic gene activity model leads to a distribution of the number of active genes - Example

In the probabilistic model, the fraction of active genes $q$ becomes a random variable, rather than being fixed, as we assumed before. We demonstrate this in an example below. The

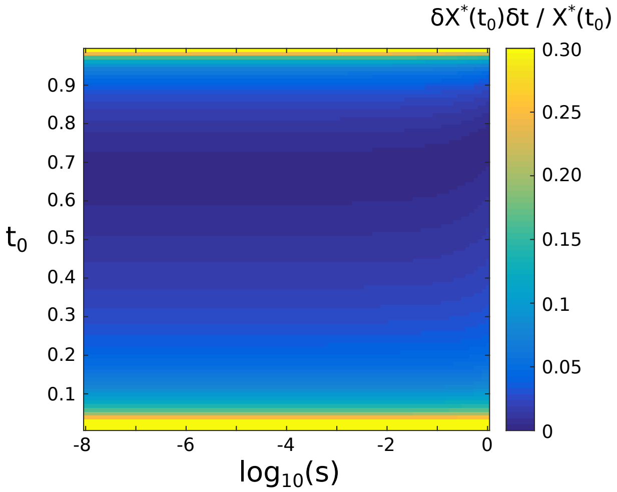

Figure S9: The relative error $\delta X^*(t_0)\delta t/X^*(t_0)$, shown in color, as a function of similarity, $s$, and the expected value of distribution t, $t_0 = \langle t \rangle$. Parameter values: $M = 2500$ and $\gamma_i = 1/2$, leading to $\delta t = 10^{-2}$. All other values of $\gamma_i$ would lead to lower values of $\delta t$ and therefore to lower values of the relative error.

crosstalk behavior in the probabilistic case can be obtained as a superposition of the relevant deterministic cases taken with their corresponding weights.

Assume we have 3 genes, active with probabilities $p_1 = 1/2$ and $p_2 = p_3 = 1/4$, correspondingly. We can then enumerate all active gene combinations and active state probabilities:

- all genes inactive: $p(\text{all inactive}) = \frac{1}{2}\left(\frac{3}{4}\right)^2 = 9/32$
- first gene active: $p(\text{gene 1 active, genes 2,3 inactive}) = \frac{1}{2}\frac{3}{4}\frac{3}{3} = 9/32$
- second gene active: $p(\text{gene 2 active, genes 1,3 inactive}) = \frac{1}{2}\frac{1}{4}\frac{3}{4} = 3/32$
- third gene active: $p(\text{gene 3 active, genes 1,2 inactive}) = \frac{1}{2}\frac{3}{4}\frac{1}{4} = 3/32$
- first&second genes active: $p(\text{genes 1,2 active, gene 3 inactive}) = \frac{1}{2}\frac{1}{4}\frac{3}{4} = 3/32$
- first&third genes active: $p(\text{genes 1,3 active, gene 2 inactive}) = \frac{1}{2}\frac{3}{4}\frac{1}{4} = 3/32$
- second&third genes active: $p(\text{genes 2,3 active, gene 1 inactive}) = \frac{1}{2}\frac{1}{4}\frac{1}{4} = 1/32$
- all genes active: $p(\text{all active}) = \frac{1}{2}\frac{1}{4}\frac{1}{4} = 1/32.$

The mean number of active genes $Q$ is then:

$$\langle Q \rangle = 0 \times p(\text{all inactive}) + 1 \times p(\text{gene 1 active, genes 2,3 inactive}) + \cdots + 3 \times p(\text{all active}) = 1. \tag{S19}$$

Therefore, in this example, on average, one gene is active.
However, for $\langle Q \rangle = 1$ and fixed $p$ (fraction of activator among the existing regulators), there are several possible $q$ values (proportion of active genes). This explains why we have a distribution of crosstalk values if genes are active with some probability.

### 8 Data-based crosstalk calculations

#### 8.1 Distribution of similarity measures for *S. cerevisiae* genes is relatively wide

To obtain similarity and crosstalk values of real organisms, one needs to take several aspects into consideration. First, the exact consensus sequences of different TFs are not known and position count matrices (PCMs) are used to infer them. Second, the length of binding sites and consensus sequences between different BSs and TFs can differ. Third, in a more realistic case, each TF can be cognate for multiple genes. All these concerns (and others, for more details, see Methods) complicate a calculation of lower bound on crosstalk in a real organism. However, there are ways to solve these issues and obtain estimations to be compared with our analytical solutions.

We define similarity between a binding site $k$ and transcription factor $l$ as $S_{kl} = \exp(-E^{kl})$, where $E^{kl}$ represents the mismatch energy of binding of the transcription factor on the binding site. The similarities between all pairs of consensus binding sites for *S. cerevisiae* are shown in Fig S10. The results are not symmetric between transcription factors and binding sites. A simple example with two transcription factors and their cognate binding sites can be presented to understand this intuitively: imagine the first transcription factor with a shorter consensus sequence while the consensus sequence of the second one is longer. Let us assume that a consensus of the shorter TF is included in the consensus of the longer TF.

The TF with the shorter consensus sequence will bind easily to the binding site of the second TF. Therefore, the similarity between the transcription factor with the shorter consensus sequence and longer binding site will be high. However, the transcription factor with the longer consensus sequence and a shorter binding site would have a lower similarity, as it is less likely that the long transcription factor binds to the short binding site.
Indeed, the matrix of pairwise similarity values is asymmetric. Clearly, we observe many vertical lines of similar value (TFs that easily bind many binding sites - yellow strips, or that are very unique to only few binding sites - blue strips), but much weaker signatures of rows (binding sites that are very similar or very dissimilar to all others). This demonstrates that the similarity value between a transcription factor and a binding site is dominated by the transcription factor properties, and much less by the binding site's. High similarity between a transcription factor and other binding sites is highly correlated with short consensus sequences of that transcription factor.

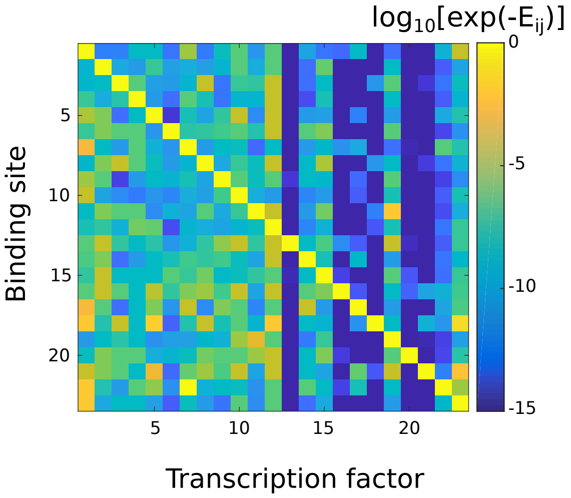

Figure S10: Similarity values between all pairs of consensus binding sites and transcription factors for *S. cerevisiae*. Columns represent TFs, rows represent binding sites.

The similarity measure of a gene $i$, $S_i$, is determined as the contribution of all non-cognate transcription factors:

$$S_i = \sum_{\text{j} \equiv \text{over all BSs}, \, j \neq i} C_j e^{-E_{ij}}, \tag{S20}$$

with $C_j$ being the concentration of TF species $j$.

##### 8.2 TFs that bind shorter DNA stretches are more promiscuous

TFs with shorter consensus BS can fit more binding sites. Subsequently, their similarity value $s_i$ is higher. This is indeed what we find in our yeast data – see Fig. S11.

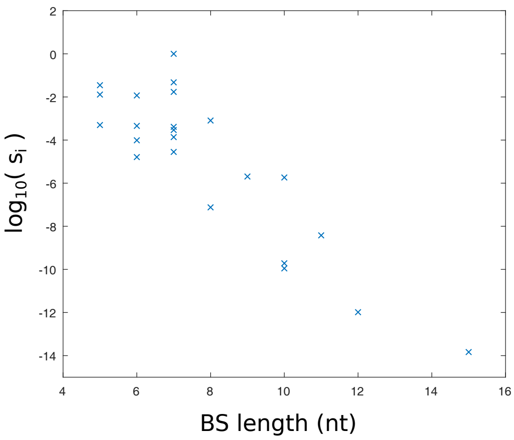

Figure S11: TFs with shorter consensus BSs tend to have higher similarity values. The data points show TF similarity values calculated for the *S. cerevisiae* dataset.

##### 8.3 Alternative calculations of similarity values and crosstalk from data

Similarity values and crosstalk of *S. cerevisiae* and other organisms were estimated [1] based on PCM data of the TFs in our previous work, but using a different computational approach. The main difference between these two calculations is that in the first approach, the similarity was calculated between a consensus sequence of a particular TF and an ensemble of binding sequences of the same length randomly drawn from a uniform distribution. It was shown there analytically that the average similarity between random sequences of length $L$ and uniform energy mismatch per position $\epsilon$ is simply $S = (1/4 + 3/4 \exp(-\epsilon))^L$, hence only this effective $\epsilon$ needs to be calculated. In contrast, in the current work, we calculate similarity between actual pairs of TF and non-cognate binding sites.

Another major difference is the calculation of mismatch energy penalties. By using the PCMs, we obtain an energy matrix which gives energy penalties for every position and every nucleotide separately. In the previous work, only an effective $\epsilon$ uniform for all positions was calculated using either of two approaches: (i) information method [10] or, (ii) pseudo-count method (also used here) [11, 12]. In the information method, the total information of the motif was calculated and then an effective $\epsilon_{\text{eff}}$ which evenly distributed this information between all $L$ positions, was calculated. The advantage of this method is the avoidance of pseudo-count usage, which could bias the results. Its major drawback is the lack of position-specific energy information which we need to calculate similarity between actual pairs of binding sites. In the pseudo-count method, a pseudo-count is added to all positions in order to avoid zero counts, which result in infinite energy penalty. While this method can provide position-specific energy values $\epsilon_j$, in the previous work, only an average of all positions $\epsilon_{\text{eff}} = \sum_j \epsilon_j$ was taken as the effective value for the similarity with respect to random sequences. In the current work, the pseudo-count method was used differently, computing the similarity measure of a gene $j$ by directly following the definition and summing the Boltzmann weights over all TFs (i.e., sum over exponents of energies, $\sum_i \exp\left(-E_{ij}\right)$). Since binding sites and TFs can have different lengths, there could be different relative positions with respect to each other, which could have different binding energies. Here, we chose the relative position with highest match (lowest energy penalty) between the binding TF $i$ and binding site $j$. States with lower energy are energetically more favorable and therefore physically more likely to occur.

This difference in estimating energy penalties leads to a different approach for computing similarity measures. In our approach, we use energy matrices to compute the energy of binding for every pair of TF-BS, i.e., $E_{ij}$.

The two distinct approaches lead to different, but similar, distributions of similarity measures for a given gene $j$, $s_j$ – Fig S12. The main difference are long tails of the current approach. The median value of the similarity with the previous approach in *S. cerevisiae* was median($s^{\text{consensus-random}}$) $= 0.8 \cdot 10^{-4}$, while in the current approach, we obtain median($s^{\text{TF-BS}}$) $= 1.4 \cdot 10^{-4}$.

Differences between $s$ values obtained in these two approaches can emanate from various sources:

- sequences of actual binding sites are not well captured by a uniform distribution because of biases in favor of AT-content. For example in our *S. cerevisiae* dataset, we have 31% of nucleotide A, 21% of C, 22% of G, and 26% of T.

- actual binding sites can vary in length; taking the relative position with best match is clearly non-random. See in Fig S12 a comparison to $s$ values calculated when the relative position is randomly selected.
- equally partitioning the total energy of the motif between all its positions consistently under-estimates the similarity.
- if actual TF-BS are considered, insufficient data can lead to biases in similarity estimates.

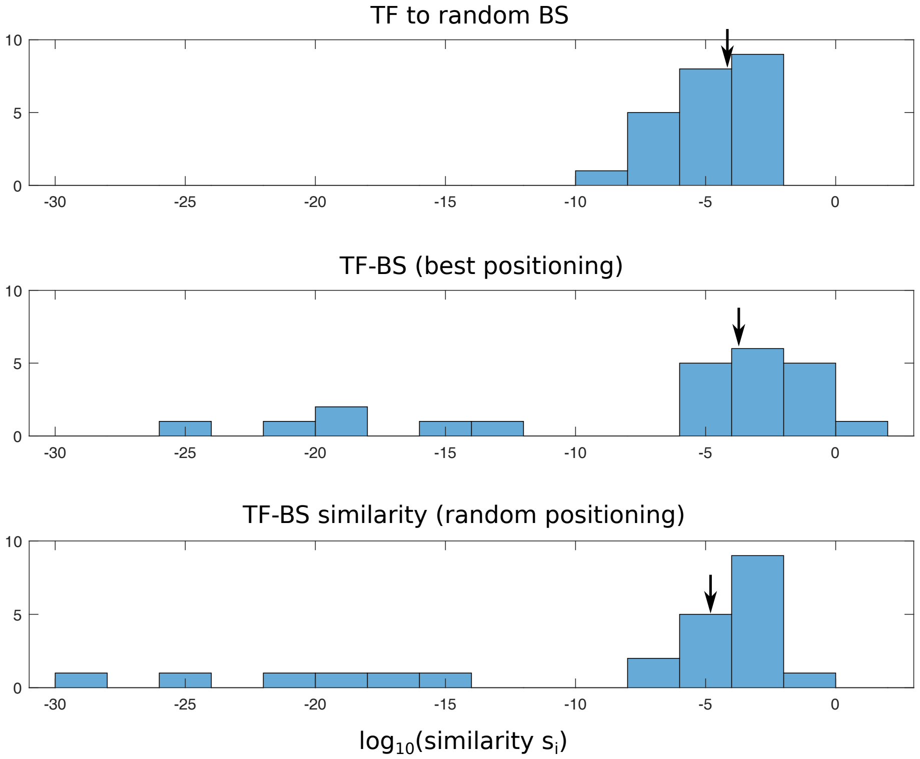

Figure S12: Comparison of distribution between the different approaches; alternative to previous work (top) versus the current work with best positioning (middle) and random positioning (bottom). Random positioning takes a random binding location instead of the one with the highest match (the distribution shown is from one representative realization). The median value (averaged over many realizations) of median$(s^{\text{TF-BS-random}}) = 0.2 \cdot 10^{-4}$.
